# Hijacking of inflammasome responses by the complement system during *Pseudomonas aeruginosa*–*Aspergillus fumigatus* sur-infection

**DOI:** 10.64898/2026.02.20.707105

**Authors:** Sandra Khau, Lucas Treps, Guy Ilango, Nicolas Riteau, Isabelle Couillin, Dieudonée Togbe, Jeanne Bigot, Viviane Balloy, Camille David, Margaux Charrier Le Blan, Delphine Fouquenet, Virginie Vasseur, Thierry Fontaine, Isabel Pappworth, Kevin Marchbank, Christophe Paget, Thomas Baranek, Elise Biquand, Clemente J Britto, Loïc Guillot, Mustapha Si-Tahar, Benoit Briard

**Affiliations:** Inserm, Centre d’Étude des Pathologies Respiratoires (CEPR), UMR1100, Tours, France; Université de Tours, Tours, France; INSERM UMR 1307, CNRS UMR 6075, Nantes Université, Université d’Angers, Nantes, F-44000, France; Experimental and Molecular Immunology and Neurogenetics, INEM UMR7355 University of Orleans and CNRS, Orleans, France; Immune Health Laboratory, Regulation of Host Responses and Immune Health, IRL2029, French National Centre for Scientific Research (CNRS) and Ribeirão Preto Medical School (FMRP) of the São Paulo University (USP), São Paulo, Brazil; Sorbonne Université, Inserm, Centre de Recherche Saint Antoine, Paris F-75012, France; Institut Pasteur, Université Paris Cité, INRAE, USC2019, Unité Biologie et Pathogénicité Fongiques, Paris, France; Complement Therapeutics Research Group, Newcastle University Translational and Clinical Research Institute, the Medical School, Newcastle-upon-Tyne, UK; National Renal Complement Therapeutics Centre, the Royal Victoria Infirmary, Newcastle-upon-Tyne, UK; Division of Pulmonary, Critical Care, and Sleep Medicine, Yale University School of Medicine, New Haven, CT, USA

**Keywords:** Inflammasome, NLRP3, C3, ITGAM, *Pseudomonas aeruginosa*, *Aspergillus fumigatus*

## Abstract

Patients with cystic fibrosis (pwCF) are highly susceptible to chronic pulmonary infections due to mutations in the CFTR gene. From early childhood, pwCF experience repeated lung infections and often develop chronic bacterial and/or fungal colonization. Among the most clinically relevant pathogens, *Pseudomonas aeruginosa* and *Aspergillus fumigatus* frequently co-infect and are associated with worse outcomes, including excessive IL-1β-driven inflammation and accelerated lung function decline. Here we investigated the mechanisms underlying inflammasome overactivation during super-infection. We found that inflammasome hyperactivation occurred across macrophage populations, was independent of exogenous priming, and required live co-infection with both pathogens. *P. aeruginosa* and *A. fumigatus* cooperatively activated the NLRP3 inflammasome, and this response required both caspase-1 and caspase-8. Unexpectedly, gasdermin D was dispensable for IL-1β release. Bacterial flagellin, type IV pili and the type III secretion system, as well as the fungal polysaccharide galactosaminogalactan (GAG), were each required for overactivation. Mechanistically, *P. aeruginosa* activated the MyD88–TLR pathway, enhancing macrophage responses and promoting ITGAM (CD11b) expression. Under fungal super-infection, macrophages secreted complement component C3, which may bound fungal surface and engaged the complement receptor C3R (CD11b/CD18). Downstream SYK and ERK signaling amplified inflammasome activation and IL-1β release. Single-cell transcriptomic analysis of pwCF broncho-alveolar lavage and lung samples supported coordinated upregulation of complement and inflammasome pathways during bacterial-fungal infection. Together, these findings identify a complement–inflammasome signaling axis that drives pathological inflammation during bacterial-fungal co-infection in airways of pwCF and may represent a therapeutic target.

## Main

Cystic fibrosis (CF) is characterized by chronic airway infection and a persistent yet ineffective inflammatory response, which remains a central challenge in patient management. This dysregulated and exaggerated inflammation drives progressive bronchiectasis and irreversible lung damage. Although inflammation is essential for pathogen control, the therapeutic goal in CF immunopathology is to mitigate excessive inflammation without compromising host defense.

Chronic infection is a major determinant of morbidity in CF, most notably involving *Pseudomonas aeruginosa*, which frequently infects and establishes persistent airway colonization early in life of patients with CF (pwCF)^1^. The pulmonary microbiota in pwCF is complex and includes diverse bacterial, viral, and fungal communities. Increasing evidence indicates that chronic polymicrobial infection represents a key pathogenic mechanism contributing to disease progression. In particular, co-colonization with *P. aeruginosa* and fungal pathogens such as *Aspergillus fumigatus* is associated with worsened pulmonary outcomes.

Chronic co-colonization with bacterial and fungal is increasingly recognized as a hallmark of advanced CF lung disease and is associated with accelerated pulmonary decline. The sequential and overlapping exposure of innate immune cells to these pathogens is characteristic of CF airways, highlighting the need to investigate immune regulation within this complex microbial environment.

Despite extensive research on CF immunopathology, the molecular mechanisms underlying the exaggerated inflammatory response remain incompletely defined. Emerging evidence suggests that bacterial and fungal co-infection induces hypersecretion of the pro-inflammatory cytokine interleukin-1β (IL-1β), which correlates with deteriorating lung function in pwCF. IL-1β is an inflammasome-dependent cytokine, produced through the activation of a cytosolic multiprotein complex that mediates caspase-1 cleavage, enabling the maturation of IL-1β and interleukine-18 (IL-18), and the induction of pyroptotic cell death.

The upstream signaling events responsible for inflammasome hyperactivation during bacterial-fungal co-infection remain poorly understood. While *P. aeruginosa* and *A. fumigatus* each activate specific inflammasome complexes when encountered individually, the sensor(s), adaptor components, and pathogen-associated molecular patterns (PAMPs) driving the synergistic hyperinflammatory response during superinfection have not yet been identified. Defining these molecular interactions and the resulting transcriptional programs will be critical to understanding how polymicrobial infections amplify inflammation and contribute to progressive lung pathology in CF. Here, we propose that complement activation hijacks inflammasome signaling in the injured lung, fueling maladaptive inflammatory loops.

## RESULTS

### Inflammasome hyperactivation in macrophages sequentially infected with *P. aeruginosa* and *A. fumigatus*

To investigate inflammasome activity and its regulation during bacterial-fungal co-infection, we established a macrophage superinfection model that reproduces the sequential exposure to *P. aeruginosa* and *A. fumigatus* observed in the airways of pwCF. Specifically, wild-type (WT) alveolar macrophages (AMs) were first infected with *P. aeruginosa* (*P.a*), followed by *A. fumigatus* (*A.f*) after resolution of the primary infection with antibiotic treatment. We then compared inflammasome activation across conditions of mono-infection (*P.a* or *A.f*) and sequential infection (*P.a→A.f*) (Fig. 1a).

**Fig. 1.**
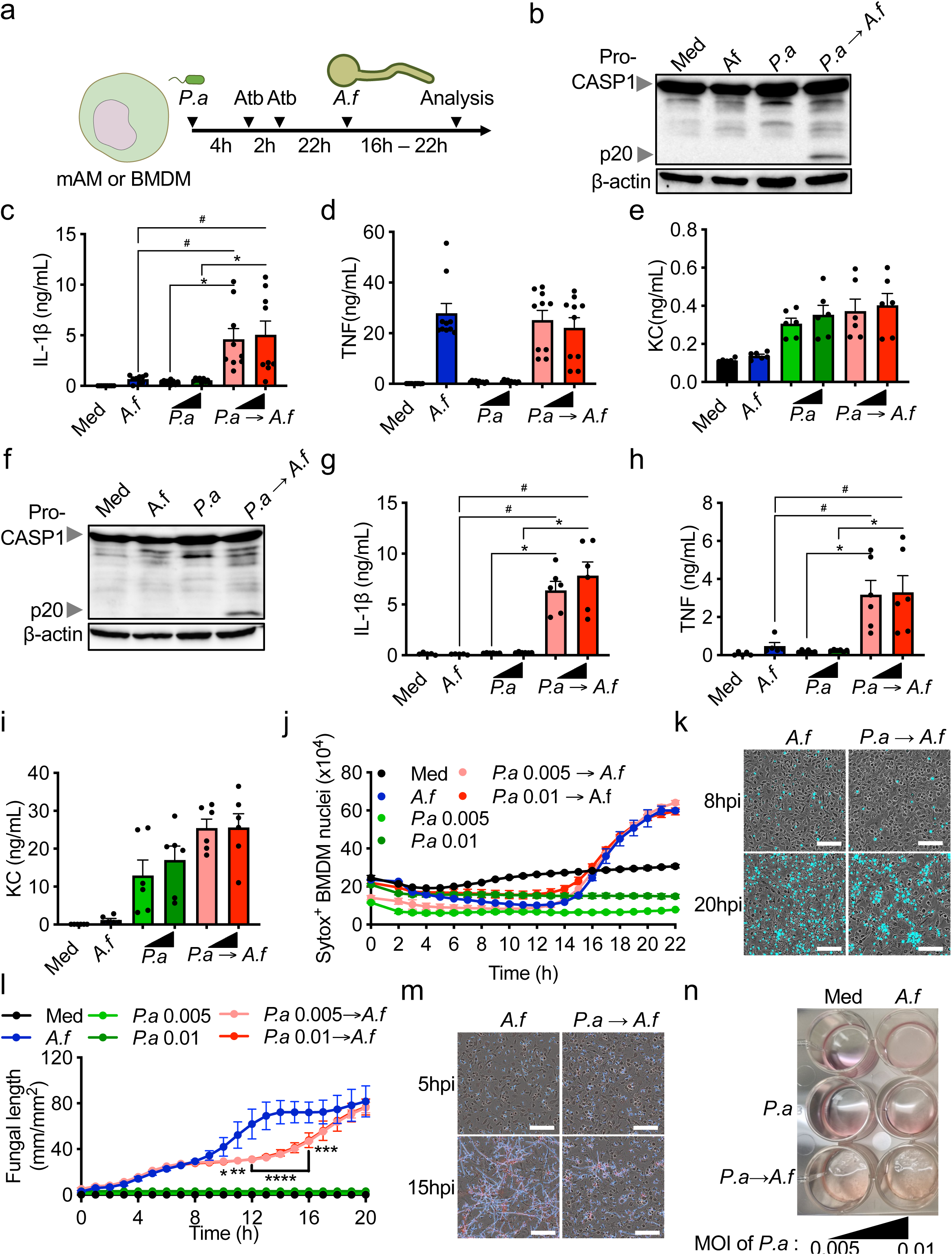
Inflammasome hyperactivation in macrophages sequentially infected with *P. aeruginosa* and *A. fumigatus*. **(a)** Experimental workflow showing primary murine alveolar macrophages (mAMs) or bone marrow-derived macrophages (BMDMs) first infected with live *P. aeruginosa* (*P.a*) for 4 hours, followed by antibiotic treatment, then challenged with *A. fumigatus* (*A.f*) conidia/hyphae, and analyzed at 16–22 hours post-superinfection. **(b)** Immunoblot of pro–caspase-1 (Pro–CASP1) and its cleaved p20 form in cell lysates from mAMs untreated (medium alone (Med)) or after infection with P.a (MOI of 0.005 or 0.01) and/or with *A.f* (MOI 5). **(c-e)** Quantification of cytokines IL-1β **(c)**, TNFα **(d)**, and KC **(e)** in supernatants from mAMs under each conditions as measured by ELISA. **(f)** Immunoblots for Pro–CASP1 and p20 in cell lysates of BMDMs from the indicated conditions. **(g-i)** BMDMs cytokine quantification of IL-1β **(g)**, TNFα **(h)**, and KC **(i)** release following *P.a* priming with increasing multiplicity of infection (MOI), and subsequent *A.f* superinfection. **(j)** Quantification of cell death, measured as the number of SytoxGreen positive (Sytox^+^) cells, in BMDMs following infection with *P.a* (MOI 0.005 or 0.01) and/or *A.f* (MOI 5). **(k)** Representative images of Sytox-stained BMDMs imaging at 8 and 20 hours post-infection (hpi) with *A.f* alone or after *P.a* primary infection “*P.a→A.f*”. Scale bars, 100µm. **(l)** Time-course analysis of fungal growth (hyphal length in mm/mm²) of *A.f* (MOI 5) after BMDMs infection, either left uninfected (medium control, Med) or co-infected with *P.a* (MOI 0.005 or 0.01). **(m)** Representative images of fungal morphology of *A.f* (Strain: DAL DSRED), highlighted in bleu, at early (5 hpi) and late (15 hpi) timepoints in BMDMs single *A.f* infection and in superinfection (MOI 5). Scale bars, 100µm. **(n)** Macroscopic images of BMDMs culture wells at endpoint (22 hpi) showing visual differences in fungal growth and cellular conditions across infection models (MOI 5 for *A.f* and MOI 0.005 or 0.01 for *P.a*). Data are representative of at least three independent experiments. Error bars indicate mean ± SEM. Statistical significance: *p<0.05, #p<0.01, ****p<0.0001, ns = not significant by ANOVA.

Sequential infection led to a pronounced increase in caspase-1 cleavage and IL-1β release compared to either mono-infection, indicating robust inflammasome activation (Fig. 1b,c). In contrast, the secretion of inflammasome-independent cytokines such as tumor necrosis factor (TNF) and keratinocyte chemoattractant (KC/CXCL1) remained unchanged under superinfection conditions (Fig. 1d,e).

To confirm these findings, we reproduced the model using WT bone marrow-derived macrophages (BMDMs). Similar to AMs, sequential infection markedly enhanced inflammasome activation in BMDMs (Fig. 1f,g). While KC release was unaffected, TNF secretion in BMDMs mirrored the pattern of IL-1β production, suggesting differential regulatory mechanisms across cytokine pathways between mAMs and BMDMs (Fig. 1h,i).

Since inflammasome activation typically induces pyroptosis, we next assessed cell death upon infection. No significant differences were observed between *A. fumigatus* mono-infected and sequentially infected BMDMs (Fig. 1j,k), suggesting that cell death in this context may occur independently of inflammasome activation. Interestingly, fungal growth was reduced during a defined growth window in *P. aeruginosa* preinfected BMDMs compared with naive macrophages (Fig. 1l–n), with altered hyphal morphology (Fig. 1n).

Together, these results demonstrate that sequential infection with *P. aeruginosa* and *A. fumigatus* drives inflammasome hyperactivation in macrophages, followed by selective modulation of cytokine responses and altered fungal growth dynamics.

### Inflammasome hyperactivation depends on macrophage potentiation by live bacterial infection

To dissect the mechanism underlying inflammasome hyperactivation, we first verified whether the bacterial infection was completely cleared before the secondary fungal challenge. Antibacterial treatment effectively eliminated *P. aeruginosa*, and no alive bacteria were detected at the time of *A. fumigatus* infection (Fig. 2a). Since inflammasome activation occurs following the secondary infection with *A. fumigatus*, the phagocytic and fungicidal capacities of macrophages upon *P. aeruginosa* may directly influence inflammasome activation. However, no significant differences were observed in either conidial phagocytosis or killing capacity (Fig. 2b,c).

**Fig. 2.**
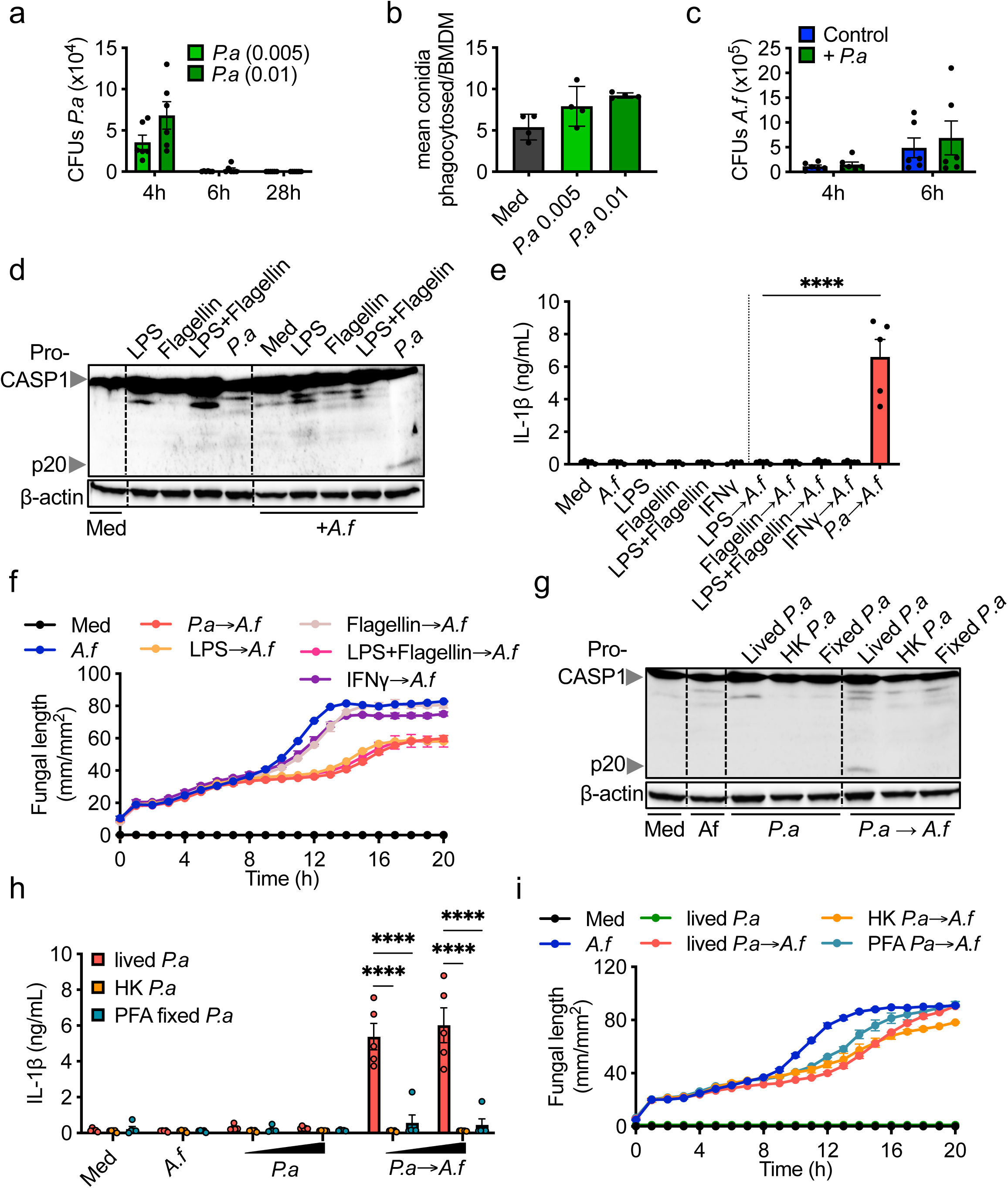
Inflammasome hyperactivation depends on macrophage potentiation by live bacterial infection. **(a)** Quantification of *P.a* colony-forming units (CFUs) in BMDMs at different time points and multiplicities of infection (MOI 0.005 or 0.01). **(b)** Mean number of *A.f* (MOI 5) conidia phagocytosed per BMDM without *P.a* priming (Med) or following priming with different doses of live *P.a* (MOI 0.005 or 0.01). **(c)** *A.f* CFUs from cultures with or without *P.a* priming at 4 and 6 hours after fungal infection, indicating enhanced fungal clearance when macrophages are pre-infected with *P.a*. **(d)** Immunoblot for pro–caspase-1 and its cleaved p20 fragment in BMDMs lysates stimulated with LPS and/or flagellin, IFNγ, or live P.a. **(e)** ELISA quantification of IL-1β secretion from BMDMs under the indicated priming (LPS alone, flagellin alone, LPS and flagellin together, or live *P.a*) and fungal challenge conditions (+*A.f*). **(f)** Time-course of *A.f* hyphal growth (length in mm/mm²) monitored in the presence of BMDMs primed with the different stimuli prior to fungal challenge. **(g)** Immunoblot for caspase-1 p20 cleavage with live *P.a* (MOI 0.01), heat-killed (HK) or PFA-fixed *P.a* (MOI 0.02) and fungal challenge (*A.f*). (h) ELISA quantification of IL-1β release upon live, HK or PFA-fixed *P.a* (with increased MOI 0.01 and 0.02) with or without second *A.f* infection (MOI 5). **(i)** Time-course quantification of *A.f* hyphal growth with BMDMs primed by live, HK (at MOI 0.02), or fixed *P.a* (MOI 0.01), in the presence or absence of *A.f* superinfection (MOI 5). Data are representative of at least three independent experiments. Error bars indicate mean ± SEM. Statistical analysis by one-way or two-way ANOVA with multiple comparisons; ****p < 0.0001.

The initial *P. aeruginosa* infection may therefore serve as a priming event, sensitizing macrophages to enhanced inflammasome activation during the subsequent fungal infection. To test this hypothesis, BMDMs were pre-stimulated with *P. aeruginosa* LPS, flagellin (alone or in combination), or IFN-γ prior to infection with *A. fumigatus*. However, none of these conditions reproduced the hyperactivation of the inflammasome observed during sequential infection with *P. aeruginosa* followed by *A. fumigatus* (Fig. 2d,e). Interestingly, fungal growth was reduced in BMDMs primed with LPS, similar to what was observed following sequential bacterial-fungal infection (Fig. 2f). This finding confirms that recognition of bacterial PAMPs can modulate macrophage responses to a secondary fungal challenge and that inflammasome activation is independent of fungal growth.

To determine whether bacterial viability was required for inflammasome hyperactivation, BMDMs were pre-exposed to fixed or heat-inactivated *P. aeruginosa* prior to *A. fumigatus* infection. Similarly to bacterial PAMPs alone, inactivated bacteria failed to induce the enhanced inflammasome activation observed with live bacterial infection (Fig. 2g,h). Similarly to the bacterial PAMPs and live bacteria, the fungal growth was reduced in BMDMs stimulated with no live *P. aeruginosa* (Fig. 2i). These results demonstrate that infection of macrophages with live *P. aeruginosa* is essential for the potentiation of the inflammasome response during subsequent *A. fumigatus* infection.

### Specific bacterial and fungal PAMPs drive inflammasome hyperactivation during sequential infection

Inflammasome activation relies on specific PAMPs recognition by cytosolic sensors that assemble multiprotein complexes to mediate caspase-1 activation and IL-1β release.

To identify the bacterial PAMPs involved, we screened a *P. aeruginosa* PAK mutant library, as this strain recapitulates the inflammasome response observed with the PAO1 strain following secondary *A. fumigatus* infection (Supplementary Fig. 1a–d). The screen revealed that flagellin (Δ*fliC*), the type III secretion system (T3SS) (Δ*pscF*), and the type IV pilus (Δ*pilQ*) were critical for inflammasome overactivation (Fig. 3a,b). To ensure these findings were not due to altered bacterial infectivity, we quantified bacterial load in BMDMs four hours post-infection and observed no significant differences among mutants (Supplementary Fig. 1e), confirming that these bacterial structures specifically contribute to macrophage response rather than infection efficiency.

**Fig. 3.**
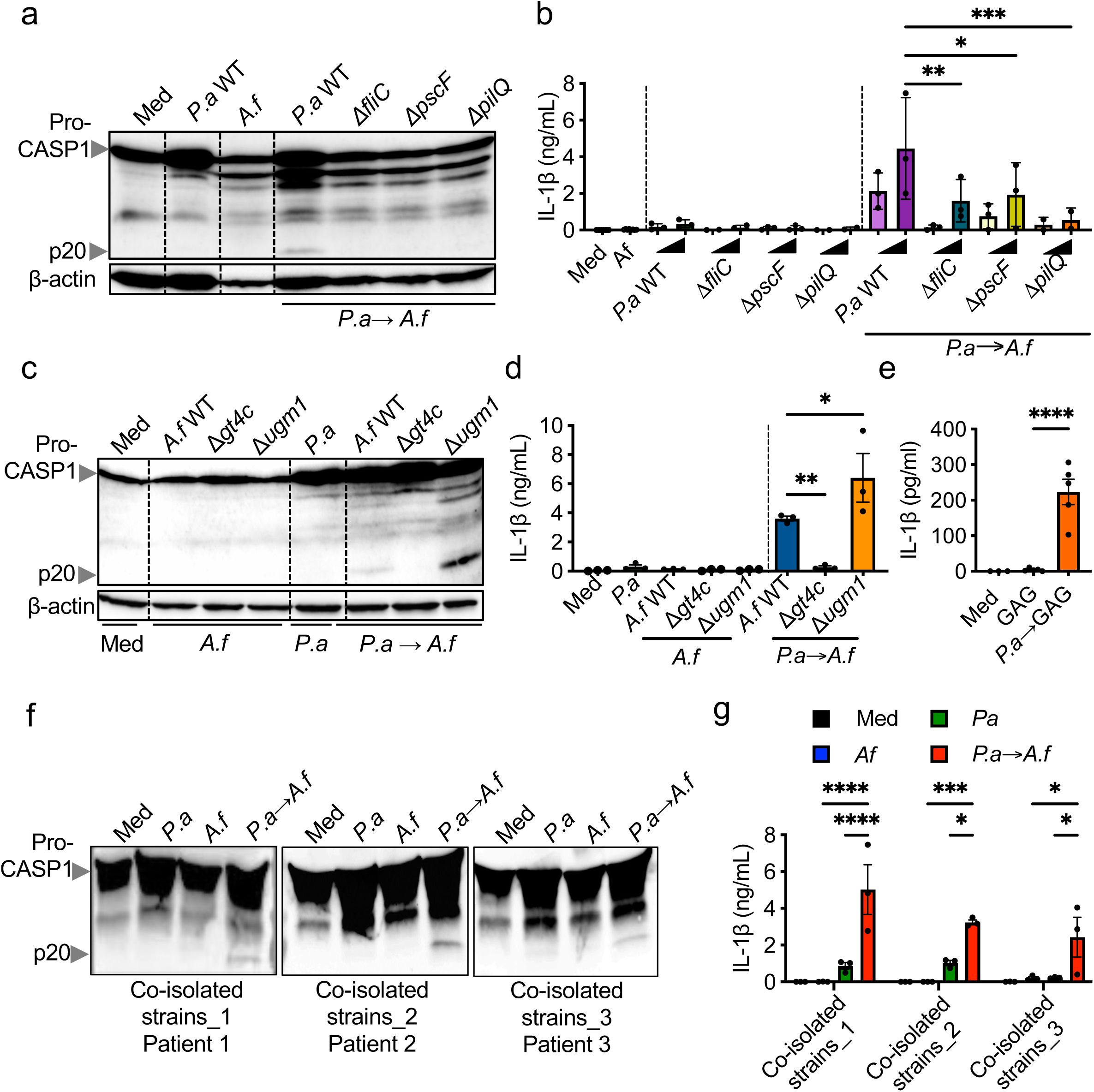
Specific bacterial and fungal PAMPs drive inflammasome hyperactivation during sequential infection. **(a)** Immunoblot for pro–caspase-1 and its cleaved p20 fragment and in BMDMs untreated (Med) or primed with wild-type (WT) or mutant *P.a* strains (Δ*fliC*, Δ*pscF*, Δ*pilQ*) at MOI of 0.01, and followed or not by superinfection with *A.f* (MOI 5). **(b)** ELISA quantification of IL-1β secretion in supernatants from BMDMs exposed to the indicated *P.a* mutants (MOI 0.005 and 0.01) and followed or not by superinfection with *A.f* (MOI 5). **(c)** Immunoblot for caspase-1 processing and **(d)** corresponding IL-1β ELISA levels in BMDMs left untreated (Med) or challenged with WT or mutant *A.f* strains (Δ*gt4c*, Δ*ugm1*) (MOI 5), following or not *P.a* priming (MOI 0.01). **(e)** IL-1β quantification in BMDMs left untreated (Med) or challenged with GAG only or first primed by *P.a* (MOI 0.01). **(f)** Immunoblots for caspase-1 cleavage and **(g)** quantification of IL-1β release in BMDMs untreated (Med) or infected with three independent sets of co-isolated clinical *P.a* (MOI 0.01) and *A.f* (MOI 5) strains from cystic fibrosis patients. Data are mean ± SEM of at least three independent experiments. Statistical significance assessed by ANOVA or t-test: *p<0.05, **p<0.01, ***p<0.001, ****p<0.0001.

On the fungal side, inflammasome activation by *A. fumigatus* is known to depend on the cell wall polysaccharide galactosaminogalactan (GAG)^2^. To evaluate its role in the context of bacterial pre-infection, we sequentially infected BMDMs with *P. aeruginosa* followed by *A. fumigatus* GAG-deficient (Δ*gt4c*) or GAG-overproducing (Δ*ugm1*) mutants. Caspase-1 cleavage and IL-1β release were markedly reduced in cells infected with the Δ*gt4c* strain compared to WT *A. fumigatus*, whereas infection with the Δ*ugm1* mutant further enhanced inflammasome activation (Fig. 3c,d). These findings confirm that GAG is a key fungal determinant of inflammasome hyperactivation in the setting of prior *P. aeruginosa* infection. Consistent with this, extracellular stimulation with purified GAG enhanced IL-1β secretion only when BMDMs were preinfected with *P. aeruginosa* (Fig. 3e), underscoring the importance of the GAG for the inflammasome activation.

Because clinical CF isolates often display strain-specific variation in PAMP composition^1^, we next examined whether the same mechanism operates in patient-derived pathogens. Sequential infection of BMDMs with co-isolated *P. aeruginosa* and *A. fumigatus s*trains from three pwCF induced robust caspase-1 cleavage and IL-1β release in each case (Fig. 3f,g), supporting the physiological relevance of this mechanism in CF. Given the complexity of the CF airway microbiota, we next assessed whether inflammasome hyperactivation is unique to *P. aeruginosa*. Sequential infection with *Escherichia coli* or *Staphylococcus aureus* likewise increased caspase-1 activation (Supplementary Fig. 1h), suggesting that this response represents a general hallmark of bacterial–fungal co-infection. Together, these results demonstrate that inflammasome hyperactivation during sequential bacterial-fungal infection depends on defined PAMPs from both alive pathogens: specifically, flagellin, T3SS, and type IV pili from *P. aeruginosa*, and GAG from *A. fumigatus*.

### Sequential infection with *P. aeruginosa* and *A. fumigatus* activates the NLRP3–ASC–caspase-1/8 inflammasome

Inflammasome activation is tightly regulated and stimulation dependent. Indeed, *P. aeruginosa* and *A. fumigatus* are each known to engage distinct inflammasome receptors. To identify the specific inflammasome receptors mediating inflammasome hyperactivation during bacterial–fungal superinfection, we infected WT, *Nlrp3^−/−^*, and *Aim2^−/−^* BMDMs with both pathogens and monitored caspase-cleavage and IL-1β release. Both caspase-1 activation and IL-1β secretion were markedly impaired only in *Nlrp3^−/−^* BMDMs, whereas TNF production remained unchanged across genotypes, as well as the impact on the *A. fumigatus* growth (Fig. 4a–d). Pharmacological inhibition of NLRP3 using MCC950 produced similar effects (Supplementary Fig. 2b–d), confirming that inflammasome activation during sequential infection is NLRP3-dependent.

**Fig. 4.**
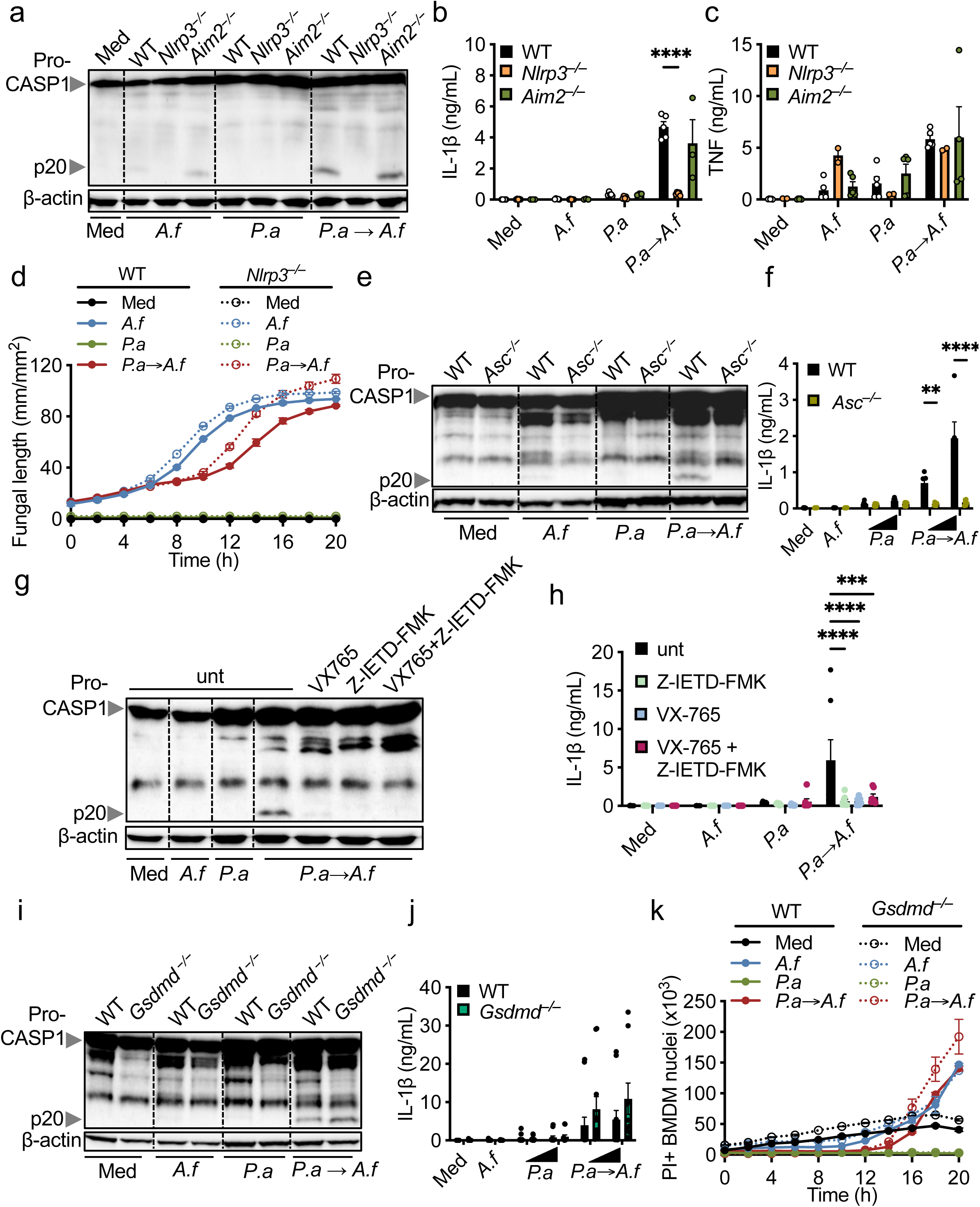
Sequential infection with *P. aeruginosa* and *A. fumigatus* activates the NLRP3–ASC–caspase-1/8 inflammasome. **(a)** Immunoblot analysis showing caspase-1 processing (pro–CASP1 and p20) and ELISA quantification of IL-1β **(b)** and TNFα **(c)** secretion in BMDMs from WT, *Nlrp3^−/−^*, and *Aim2^−/−^* mice left uninfected (Med) or after infection with *P.a* (MOI 0.005 or 0.01), and/or *A.f* (MOI 5). **(d)** Time-course of fungal hyphal growth (length/mm²) in WT and *Nlrp3^−/−^*BMDMs left uninfected (Med) or after infection with *P.a* (MOI 0.005 or 0.01), and/or *A.f* (MOI 5). **(e)** Immunoblot analysis showing caspase-1 processing (pro–CASP1 and p20) and **(f)** ELISA quantification of IL-1β secretion in BMDMs from WT, and *Asc^−/−^* mice left uninfected (Med) or after infection with *P.a* (MOI 0.005 or 0.01), and/or *A.f* (MOI 5). **(g)** Immunoblot for caspase-1 in BMDMs treated with caspase-1 inhibitor (VX765), caspase-8 inhibitor (Z-IETD-FMK), or both (VX765+Z-IETD-FMK), under single and sequential infections. **(h)** IL-1β measurement by ELISA in BMDMs treated with caspase-8 inhibitor Z-IETD-FMK, caspase-1 inhibitor VX765, their combination, left uninfected (Med) or after infection with *P.a* (MOI 0.005 and 0.01), and/or *A.f* (MOI 5). **(i)** Immunoblot analysis showing caspase-1 processing (pro–CASP1 and p20) in BMDMs from WT and *Gsdmd^−/−^* mice left uninfected (Med) or after infection with *P.a* (MOI 0.005 or 0.01), and/or *A.f* (MOI 5) and **(j)** corresponding IL-1β quantification for *Gsdmd^−/−^***. (k)** Quantification of cell death, measured as the number of propidium iodide-positive (PI^+^) cells, in *Gsdmd^−/−^* BMDMs following infection with *P.a* (MOI 0.01) and/or *A.f* (MOI 5). Data are from at least three independent experiments and presented as mean ± SEM. Statistical significance determined by ANOVA; *p<0.05, **p<0.01, ****p<0.0001.

Consistent with previous observations, *A. fumigatus* growth and macrophage cell death were unaffected by inflammasome deficiency, indicating that these processes occur independently of inflammasome activation (Fig. 4d; Supplementary Fig. 2a). Because *P. aeruginosa* can trigger inflammasome activation independently of the ASC (Apoptosis-associated Speck-like protein containing a CARD) adaptor protein^3^, we next examined its requirement. Infection of *Asc^−/−^* BMDMs revealed that caspase-1 cleavage and IL-1β release were entirely dependent on ASC (Fig. 4e–h). Then infection of *Casp1^−/−^*BMDMs confirmed the requirement of caspase-1 for IL-1β release (Supplementary Fig. 2f–h), demonstrating that the NLRP3–ASC–caspase-1 axis mediates IL-1β response.

Although caspase-1 is the canonical protease for IL-1β maturation, fungal inflammasome activation has also been linked to caspase-8 activity^4,5^. To assess its contribution, WT BMDMs were treated with inhibitors targeting caspase-1 (VX-765), caspase-8 (z-IETD-fmk), or all caspases (z-VAD-fmk). Inhibition of either caspase-1 or caspase-8 reduced caspase-1 cleavage and IL-1β release, indicating that both proteases contribute to inflammasome activation in this context (Fig. 4g,h). Infection of *Casp1^−/−^* BMDMs confirmed the requirement of caspase-1 for IL-1β release (Supplementary Fig. 2f–h).

Gasdermin D (GSDMD) is the executioner of pyroptosis downstream of inflammasome activation and has been reported to amplify NLRP3 signaling^6^. To determine its role during bacterial–fungal superinfection, we infected *Gsdmd^−/−^* BMDMs and assessed inflammasome activation. Caspase-1 cleavage, IL-1β release, and macrophage cell death were unaffected by the loss of GSDMD (Fig. 4i–k), demonstrating that inflammasome activation and cell death in this model are GSDMD-independent. Consistent with our earlier findings, the reduction in *A. fumigatus* growth in *P. aeruginosa*-preinfected BMDMs, as well as cell death induction by *A. fumigatus*, occurred independently of inflammasome activation (Fig. 4d,k; Supplementary Fig. 2a,e,h,j).

Collectively, our results identify the NLRP3–ASC–caspase-1/8 axis as the central platform mediating inflammasome hyperactivation during bacterial-fungal superinfection, uncoupled from GSDMD-driven pyroptotic cell death.

### P. aeruginosa potentiates macrophage inflammasome activation by reprogramming transcriptional and signaling responses to A. fumigatus

Inflammasome activation is controlled by complex signaling networks that vary depending on the nature and pathogen exposure. To investigate the mechanisms underlying inflammasome hyperactivation during sequential infection, we performed transcriptomic profiling of BMDMs infected or not with *P. aeruginosa*, followed by secondary infection with *A. fumigatus* for 4 or 8 hours (corresponding to 32- and 36-hour time points; see Fig. 1a).

Principal component analysis (PCA) revealed clear clustering by infection status and time point (Supplementary Fig. 3a,b). Interestingly at the pre-fungal stage (28 h), *P. aeruginosa* infection alone did not significantly alter global transcriptional variance or gene expression compared with uninfected controls. In contrast, after *A. fumigatus* challenge, *P. aeruginosa*-infected macrophages displayed distinct clustering from naive cells, indicating that prior bacterial infection profoundly reshaped the transcriptional response to fungal exposure (Supplementary Fig. 3a,b).

Differential expression analysis identified a set of genes significantly modulated in *P. aeruginosa*-infected macrophages upon *A. fumigatus* superinfection, when compared to *A. fumigatus* mono-infection (|log₂ (fold change)| > 1; FDR < 0.05; Fig. 5a). As expected, *Il1b* was strongly upregulated, along with interferon-stimulated genes (ISGs) involved in antimicrobial responses (*Irf7*, *Ifitm2*, *Ifitm3*, *Isg15*, *Oas1a*), and innate immune receptors and regulators associated with antifungal defense (*Clec4e*, *Clec4n*, *Malt1*, *Hif1a*). Gene ontology (GO) enrichment analysis showed significant overrepresentation of terms related to inflammatory and innate immune responses, including “response to bacterium,” “defense response,” and “cytokine-mediated signaling” (Fig. 5b). The 30 most upregulated genes confirmed *Il1b* as one of the most responsive cytokines to prior *P. aeruginosa* exposure (Fig. 5c). This transcriptional pattern remained consistent across time points, indicating a stable effect on macrophage activation dynamics.

**Fig. 5.**
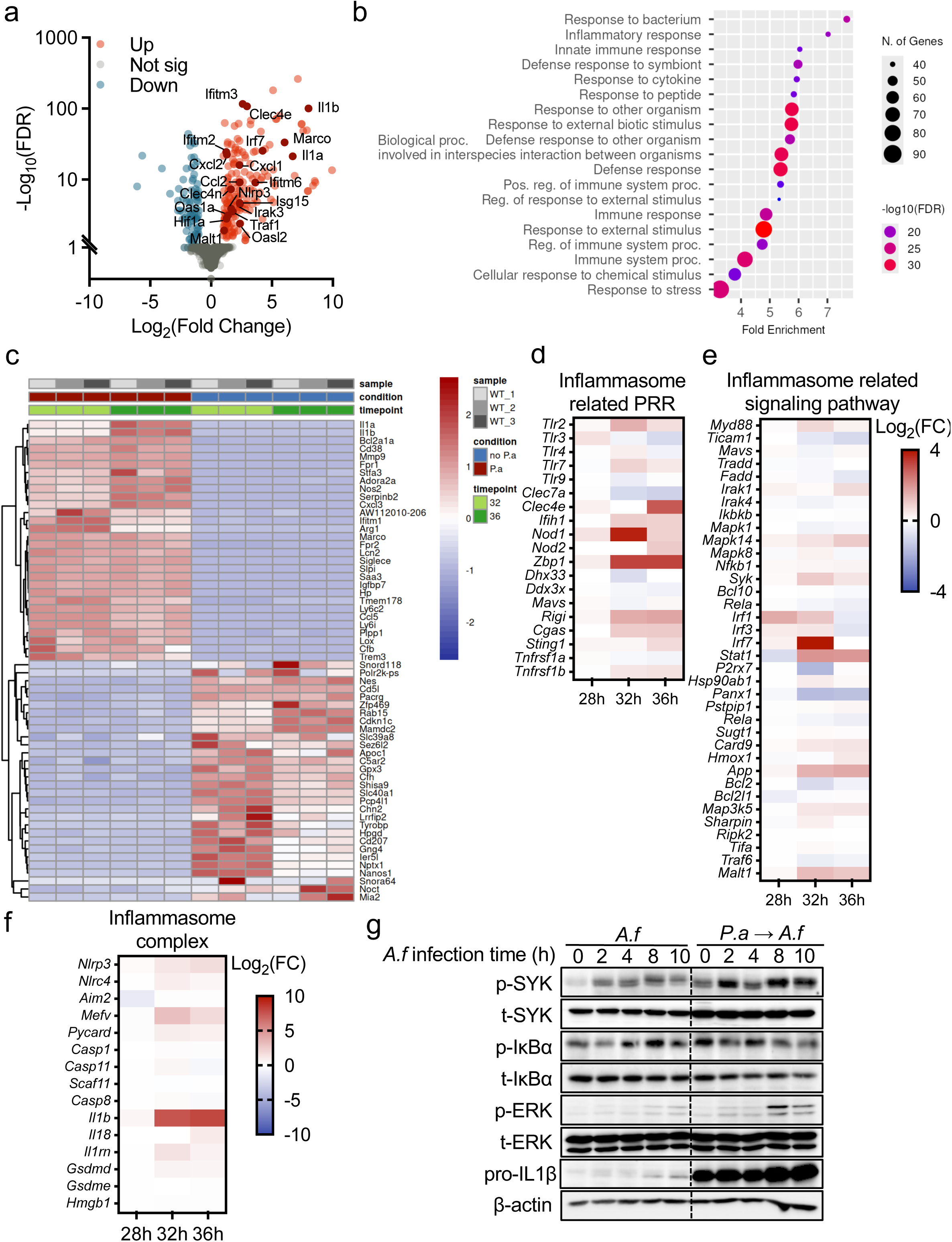
*P. aeruginosa* potentiates macrophage inflammasome activation by reprogramming transcriptional and signaling responses to *A. fumigatus*. **(a)** Volcano plot illustrating differentially expressed genes (DEGs) in BMDMs comparing *P.a*→*A.f* superinfection *vs. A.f* single infection at 36 h time point. Upregulated (red), downregulated (blue), and non-significant (grey) genes are indicated; key inflammatory genes are labeled. **(b)** Gene ontology (GO) enrichment analysis of significantly upregulated transcripts following superinfection, highlighting overrepresented biological process categories related to immune and inflammatory responses in BMDMs sequentially infected by *P.a*→*A.f*. **(c)** Heatmap showing the top 30 up- and downregulated genes in WT BMDMs across samples, stratified by condition (*P. aeruginosa* pre-exposure (*P.a*) vs no *P.a*) and time point (32 h = 4 h post–*A. fumigatus* infection; 36 h = 8 h post–*A. fumigatus* infection). **(d–f)** Heatmaps showing log2 fold change (log2FC) in WT BMDMs with *P. aeruginosa* pre-exposure (*P.a*) *vs*. no *P.a* at baseline before *A. fumigatus* infection (28 h), and at 4 h (32 h) and 8 h (36 h) post–*A. fumigatus* infection. Heatmaps highlight genes encoding **(d)** inflammasome-related PRRs, **(e)** inflammasome-associated signaling pathways, and **(f)** core inflammasome components. **(g)** Immunoblot analysis of phosphorylated and total SYK, IκBα, and ERK, and of pro-IL-1β, in BMDM lysates collected over a time course (0, 2, 4, 8, 10 hours) post-*A.f* (MOI 5) alone or post *A.f* infection in BMDMs already primed by *P.a* (MOI 0.01) *P.a*→*A.f*. β-actin is shown as a loading control.

To determine whether this effect reflected changes in inflammasome pathway gene expression, we examined transcripts specifically associated with inflammasome signaling (Supplementary Table 1). Notably, genes encoding canonical inflammasome sensors (PRRs), adaptor proteins, and regulators showed no clear differences between *P. aeruginosa*-infected macrophages upon *A. fumigatus* superinfection, when compared to *A. fumigatus* mono-infection (Fig. 5d-f). These findings suggest that inflammasome hyperactivation arises from altered upstream signaling mechanism rather than transcriptional changes in inflammasome components themselves.

We next assessed key signaling pathways known to regulate inflammasome priming and activation in response to *A. fumigatus*. In agreement with transcriptomic data, pro-IL-1β protein levels were markedly increased in BMDMs sequentially infected with *P. aeruginosa* and *A. fumigatus* compared to mono-infected cells (Fig. 5g). Phosphorylation of IκBα was comparable between conditions, indicating similar NF-κB activation (Fig. 5g). However, phosphorylation of spleen tyrosine kinase (SYK) and extracellular signal-regulated kinases 1/2 (ERK1/2) was substantially enhanced in macrophaes previously infected with *P. aeruginosa* upon *A. fumigatus* infection (Fig. 5g). While total ERK1/2 protein levels were unchanged, SYK protein abundance increased following *P. aeruginosa infection*, suggesting that bacterial priming promotes the accumulation and hyperactivation of SYK, which in turn amplifies MAPK signaling during subsequent fungal exposure.

Together, these results demonstrate that primary infection with *P. aeruginosa* functionally potentiates macrophages for enhanced inflammasome activation in response to *A. fumigatus*. This priming establishes a distinct transcriptional and signaling signature characterized by increased expression of inflammatory and interferon-stimulated genes and potentiation of SYK–ERK-dependent pathways, providing a mechanistic basis for the hyperinflammatory phenotype observed during bacterial-fungal superinfection.

### MyD88 signaling is a key regulator of inflammasome hyperactivation during sequential infection with *P. aeruginosa* and *A. fumigatus*

Because transcriptional and translational changes do not always reflect the functional impact of upstream signaling, we investigated the contribution of major innate immune pathways to inflammasome regulation in our sequential infection model. We infected WT, *Tnf^−/−^*, *Ifnar1^−/−^*, and *Myd88^−/−^* BMDMs sequentially with *P. aeruginosa* and *A. fumigatus* and assessed caspase-1 activation and IL-1β secretion. Deletion of *Ifnar1* or *Tnf* did notreduce caspase-1 cleavage or IL-1β release, indicating that type I interferon and TNF signaling are not directly involved in inflammasome activation under these conditions (Supplementary Fig. 4). In contrast, *Myd88^−/−^*BMDMs displayed markedly reduced caspase-1 activation and IL-1β release compared with WT cells (Fig. 6a,b), identifying MyD88 as a critical mediator of inflammasome hyperactivation during bacterial–fungal superinfection.

**Fig. 6.**
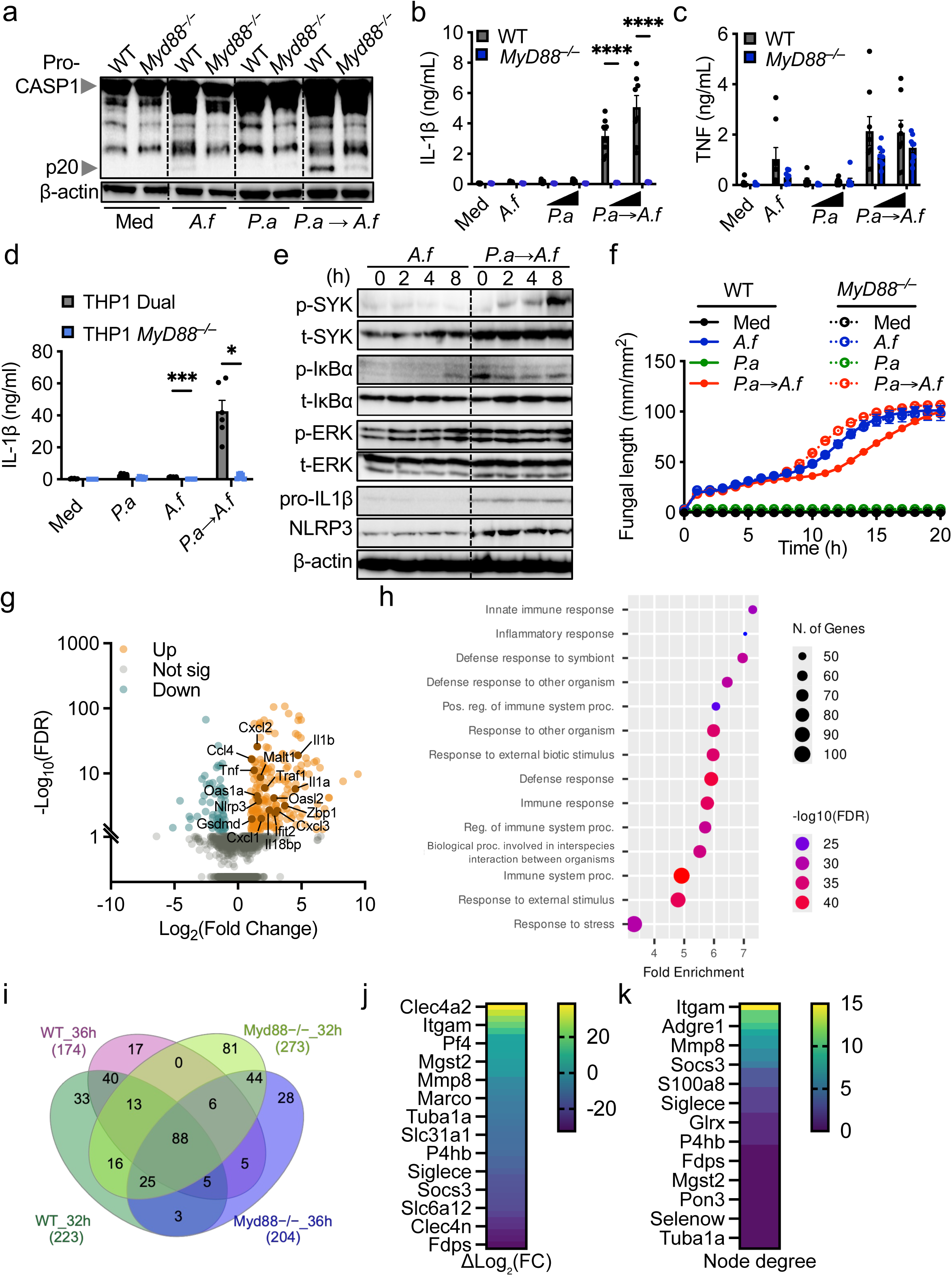
MyD88 signaling is a key regulator of inflammasome hyperactivation during sequential infection with *P. aeruginosa* and *A. fumigatus*. **(a)** Immunoblot analysis showing caspase-1 processing (pro–CASP1 and p20 cleavage) and ELISA quantification of IL-1β **(b)**, TNFα **(c)** secretion in BMDMs from WT and *Myd88^−/−^* mice left uninfected (Med) or after infection with *P.a* (MOI 0.005 or 0.01), and/or *A.f* (MOI 5). (a) Data are means ± SEM (****p<0.0001 by ANOVA). **(d)** Quantification of IL-1β levels in supernatants from WT THP-1 Dual and *MyD88*^−/−^ knockout THP-1 macrophages after left uninfected (Med), infected by *P.a* alone (MOI 0.01), *A.f* alone (MOI 5) or sequential infection (*P.a→A.f*). Results are presented as means ± SEM (*p<0.05, ***p<0.001 by ANOVA). **(e)** Immunoblot of phospho-SYK (p-SYK), total SYK, phospho-IκBα (p-IκBα), total IκBα, phospho-ERK (p-ERK), total ERK, pro-IL-1β, NLRP3, and β-actin over indicated time points after *A.f* single infection (MOI 5) or *P.a→A.f* superinfection (*P.a* MOI 0.01 and *A.f* MOI 5). **(f)** Quantification of *A.f* hyphal growth (length/mm² over time) in WT and *Myd88*^−/−^ BMDMs left uninfected (Med) or after infection with *P.a* (MOI 0.01), and/or *A.f* (MOI 5). **(g)** Volcano plot of RNA-seq analysis comparing WT and *Myd88*^−/−^ BMDMs following *P.a→A.f* superinfection, highlighting upregulated (orange) and downregulated (blue) genes. **(h)** Gene ontology (GO) enrichment for differentially expressed genes, showing biological processes with greatest fold enrichment in *Myd88^−/−^* BMDMs sequentially infected by *P.a→A.f*. **(i)** Venn diagram showing overlap of differentially expressed genes among WT and *Myd88^−/−^* BMDMs across the given timepoints and conditions. **(j)** Heatmap of top upregulated genes based on fold change (by log2 fold change, ΔLog2FC) involved in immune response. **(k)** Network node degree heatmap of top genes, underscoring interaction strength and centrality in the regulatory network.

Analysis of inflammasome-independent cytokines revealed distinct regulatory patterns: secretion of KC was completely abrogated in *Myd88^−/−^*BMDMs, whereas TNF release remained unaffected (Fig. 6c; Supplementary Fig. 5a). These findings indicate that MyD88 contributes to macrophage priming but that different cytokine pathways diverge downstream of this adaptor. Consistent with these results, MyD88 dependence was also observed in human THP-1-derived macrophages (Fig. 6d; Supplementary Fig. 5c), highlighting the relevance of this pathway in CF-related inflammation across species.

At the protein level, *Myd88^−/−^* BMDMs infected sequentially with *P. aeruginosa* and *A. fumigatus* showed impaired inflammasome activation but increased expression of pro-IL-1β and NLRP3 compared with cells infected with *A. fumigatus* alone (Fig. 6e). SYK phosphorylation was also elevated, whereas ERK1/2 and NF-κB phosphorylation appeared unaltered, suggesting that MyD88 selectively regulates upstream signaling modules involved in inflammasome activation (Fig. 6e).

Because MyD88 serves as a central adaptor for both Toll-like receptor (TLR) and IL-1 receptor (IL-1R) pathways, we next sought to distinguish their respective contributions. Sequential infection of *Il1r1^−/−^*BMDMs showed no reduction in caspase-1 activation, IL-1β or TNF secretion, cell death, or fungal growth compared with WT controls (Supplementary Fig. 5c–g). These results indicate that the role of MyD88 in inflammasome activation is primarily mediated through TLR signaling rather than IL-1R feedback loop. In agreement, the reduction in *A. fumigatus* growth observed in *P. aeruginosa*-preinfected WT macrophages was lost in *Myd88^−/−^* BMDMs (Fig. 6f), reinforcing the importance of TLR–MyD88-dependent bacterial recognition in establishing macrophage potentiation in response to *A. fumigatus* (see Fig. 2f).

To further elucidate how MyD88 regulates inflammasome activation during sequential infection, we performed transcriptomic analysis of *Myd88^−/−^* BMDMs infected or not with *P. aeruginosa*, followed by secondary infection with *A. fumigatus* for 4 or 8 hours (corresponding to 32- and 36-hour time points; see Fig. 1a). PCA revealed distinct clustering by infection status and time point (Supplementary Fig. 6a,b). Moreover, *P. aeruginosa* pre-infection induces transcriptional divergence upon *A. fumigatus* challenge, suggesting also transcriptional reprogramming in the absence of MyD88. Indeed, differential expression analysis identified genes significantly modulated in *Myd88^−/−^*BMDMs upon sequential infection (|log₂ fold change| > 1; FDR < 0.05; Fig. 6g). Unexpectedly, *Il1b* remained upregulated in *Myd88^−/−^* BMDMs *P. aeruginosa* and *A. fumigatus* superinfected compared with *A. fumigatus* monoinfection, along with other inflammasome-related genes (*Nlrp3*, *Il18bp*, *Gsdmd*), ISGs (*Ifit2*, *Zbp1*, *Oas1a*, *Oasl2*), and inflammatory cytokines (*Tnf*, *Ccl4*, *Cxcl1*, *Cxcl2*, *Cxcl3*, *Il1a*) as well as the immune regulator *Malt1* during fungal infection (Fig. 6g, supplementary Fig. 6c). GO enrichment analysis showed overrepresentation of terms related to innate and inflammatory responses, comparable to WT BMDMs (Fig. 6h and 5b). GO enrichment analysis showed overrepresentation of terms related to innate and inflammatory responses, comparable to WT BMDMs (Fig. 6h and 5b). Similarly, the heatmap of top 30 genes up and down regulated in *MyD88^−/−^* prior infected or not by *P.* show the similar signature, with the presence of key up regulated genes as *Il1b*, *Il1a, Nos2, Mmp9*, *Marco* and other in BMDMs infected by *P. aeruginosa* then *A. fumgiatus* (Supplementary Fig. 6c). All of these results indicates that loss of MyD88 does not abolish global immune activation but may alter specific regulatory pathways controlling inflammasome function.

To pinpoint genes potentially responsible for impaired inflammasome activation in *Myd88^−/−^* macrophages, we compared the lists of significantly upregulated genes between WT and *Myd88^−/−^*BMDMs after sequential infection. Forty genes were uniquely upregulated in WT cells across both fungal infection time points (Fig. 6i). We ranked these genes by their fold-change increase over time and identified twelve showing progressive upregulation during infection (Fig. 6j). Protein-interaction network analysis (STRING) of the forty WT-specific candidates revealed several highly connected clusters (Supplementary Fig. 6d), with ITGAM displaying the highest node degree (Fig. 6k).

Together, these results identify MyD88 as a central regulator of inflammasome activation during sequential *P. aeruginosa*–*A. fumigatus* infection and pinpoint ITGAM as a key downstream component potentially mediating MyD88-dependent inflammasome overactivation.

### ITGAM–C3 axis drives inflammasome hyperactivation during sequential infection with *P. aeruginosa* and *A. fumigatus*

ITGAM (also named CD11b) serves as a marker of macrophage activation and differentiation. Following *P. aeruginosa* infection and prior to *A. fumigatus* exposure, ITGAM expression was markedly increased compared with uninfected control BMDMs (Fig. 7a). To assess whether ITGAM contributes to inflammasome activation during bacterial–fungal superinfection, we treated BMDMs with a neutralizing antibody against ITGAM and evaluated caspase 1 cleavage and IL-1β release. Both caspase-1 activation and IL-1β secretion were strongly reduced upon ITGAM blocking compared to controls (Fig. 7b,c), identifying ITGAM as a critical mediator of inflammasome hyperactivation during sequential infection.

**Fig. 7.**
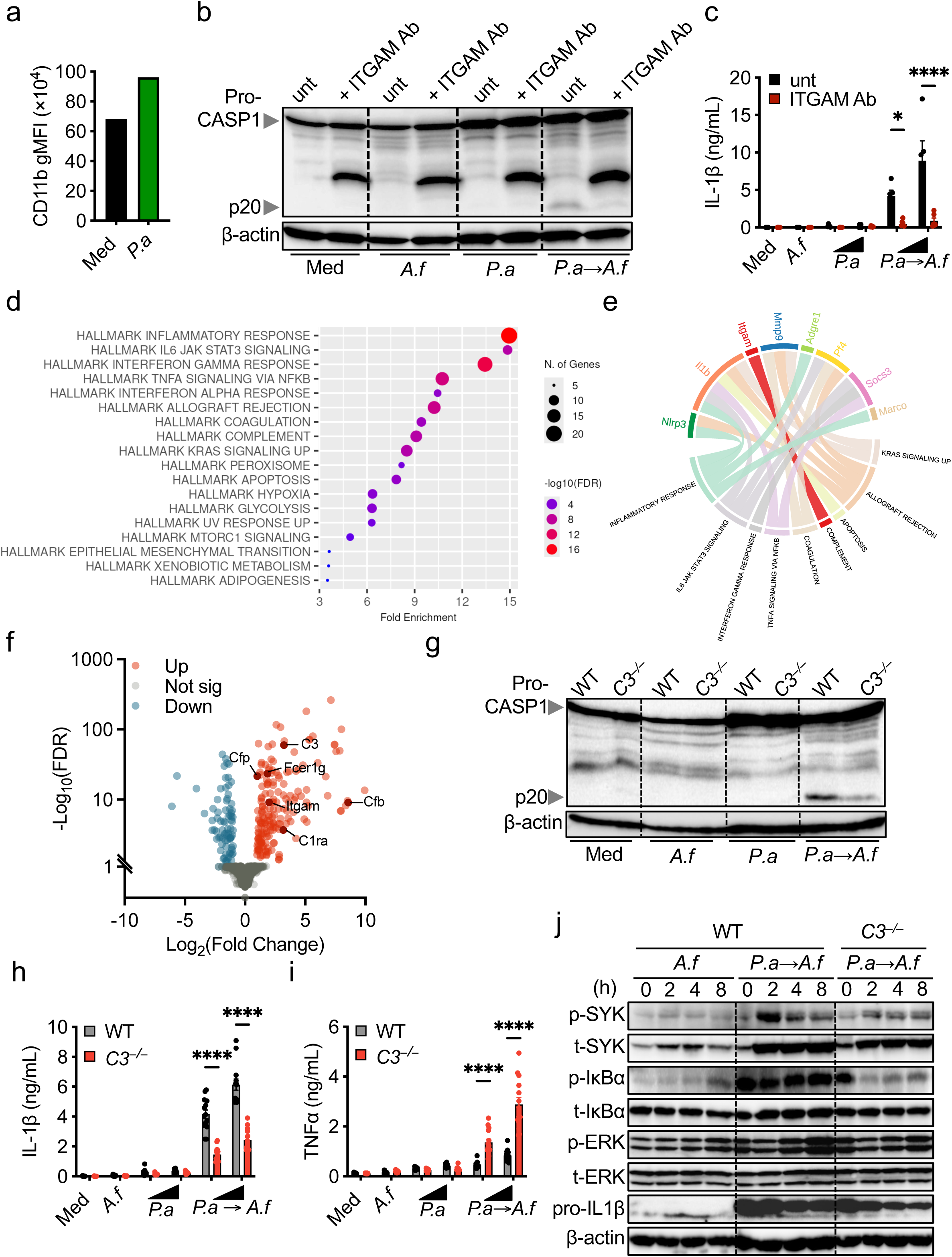
CD11b integrin participates in the inflammasome overactivation during *P.a*→*A.f* superinfection. **(a)** Quantification of CD11b surface expression (general mean fluorescence intensity, gMFI) in BMDMs following incubation with medium (Med) or *P.a,* assessed by flow cytometry. **(b)** Immunoblot showing caspase-1 activation (pro-caspase-1 and p20 cleavage fragment) and **(c)** corresponding IL-1β levels in WT BMDMs untreated (Med), or infected with *A.f* alone, *P.a* alone or sequential infection (*P.a*→*A.f*), with or without anti-ITGAM antibody blockade. **(d)** Dot plot of hallmark pathway enrichment for differentially expressed genes, showing fold enrichment, number of genes, and statistical significance by -log10(FDR). **(e)** Chord diagram of network connectivity among genes (top part of the diagram) and the pathways (bottom part) in immune and inflammatory responses. **(f)** Volcano plot showing differentially expressed genes in *P. aeruginosa*–infected WT BMDMs upon *A. fumigatus* infection, with significantly upregulated (red), downregulated (blue) and non-significant (grey) genes. **(g)** Immunoblot analysis showing caspase-1 processing (pro–CASP1 and p20 cleavage) and ELISA quantification of IL-1β **(h)** and TNFα **(i)** in BMDMs from WT and *C3^−/−^* mice left uninfected (Med) or after infection with *P.a* (MOI 0.005 or 0.01), and/or *A.f (*MOI 5). **(j)** Immunoblot time course investigating protein phosphorylation of SYK, ERK, IκBα, and pro-IL1β processing in WT and *C3^−/−^* BMDMs after *A.f* or sequential *P.a*→*A.f* infection for the indicated hours post-infection (hpi). β-actin is shown as a loading control.

To gain insight into the molecular pathways downstream of ITGAM, we performed Hallmark gene set enrichment analysis (GSEA) on transcriptomic data from WT BMDMs infected with *A. fumigatus* with or without prior *P. aeruginosa* exposure. Enrichment analysis confirmed the upregulation of pathways associated with the inflammatory response and JAK–STAT signaling, including those induced by IL-6 and type I and type II interferons (Fig. 7d). To further explore ITGAM-associated processes, we examined pathway associations between *Itgam*, inflammasome-related genes, and other key regulators identified previously (Fig. 6j–k). Notably, *Itgam* was uniquely enriched within the complement activation hallmark (Fig. 7e). Indeed, ITGAM is an integrin subunit that forms the complement receptor 3 (CR3) complex with ITGB2 (also named CD18).

Then, we analyzed complement-related genes that were differentially expressed in *P. aeruginosa*–infected BMDMs upon *A. fumigatus* infection (|log₂ fold change| > 1; FDR < 0.05). This analysis revealed strong upregulation of *C3* among the top differentially expressed genes (Fig. 7f; Supplementary Fig. 7), together with components of the alternative complement pathway, including *Cfp, Cfb*, *C1qa*, *C1ra* and *Itgam*. These findings highlight activation of the complement system, which has previously been linked to inflammasome regulation^7^.

To test whether complement signaling contributes functionally to inflammasome activation in this context, we performed sequential infections in WT and *C3^−/−^*BMDMs. In the absence of C3, caspase 1 cleavage and IL-1β release were markedly reduced compared with WT BMDMs, whereas TNF secretion was not diminished and even increased (Fig. 7g–i). Moreover, phosphorylation of SYK and NF-κB, previously enhanced in WT cells during sequential infection, was abolished in *C3^−/−^* BMDMs (Fig. 7j), indicating that complement signaling via the ITGAM–C3 axis regulates these upstream pathways.

Together, these results demonstrate that the ITGAM–C3 axis controls macrophage inflammasome activation during bacterial–fungal superinfection by modulating cytoplasmic signaling cascades, including SYK and NF-κB pathways.

### Single-cell transcriptomic profiling of pwCF identifies macrophage subsets expressing inflammasome and complement signatures

Having established that complement modulates inflammasome activation during bacterial–fungal superinfection, we next sought to evaluate whether these pathways are coordinately engaged in airways of pwCF. We analyzed publicly-available single-cell RNA sequencing (scRNAseq) datasets on sputum and lung biopsy samples obtained from pwCF, stratified by *P. aeruginosa* and/or *A. fumigatus* infection status (Supplementary Table 2). Approximately half of the individuals were clinically stable at sampling, whereas the remainders were sampled during infectious exacerbations. All pwCF carried at least one F508del *CFTR* allele, with most being homozygous (Supplementary Table 2).

Integration and unsupervised clustering identified 10 major cell populations and 7 immune cell populations across sputum and lung tissue (Fig. 8a, Supplementary Fig. 8a). Inflammasome-associated gene expression was most pronounced within macrophage, dendritic cell (DC), and neutrophil clusters (Fig. 8b-c and Supplementary Fig. 8b-c), whereas complement-associated gene expression was associated with macrophages, DC, neutrophils, epithelial, stromal and endothelial clusters (Fig. 8b-c and Supplementary Fig. 8b-c).

**Fig. 8.**
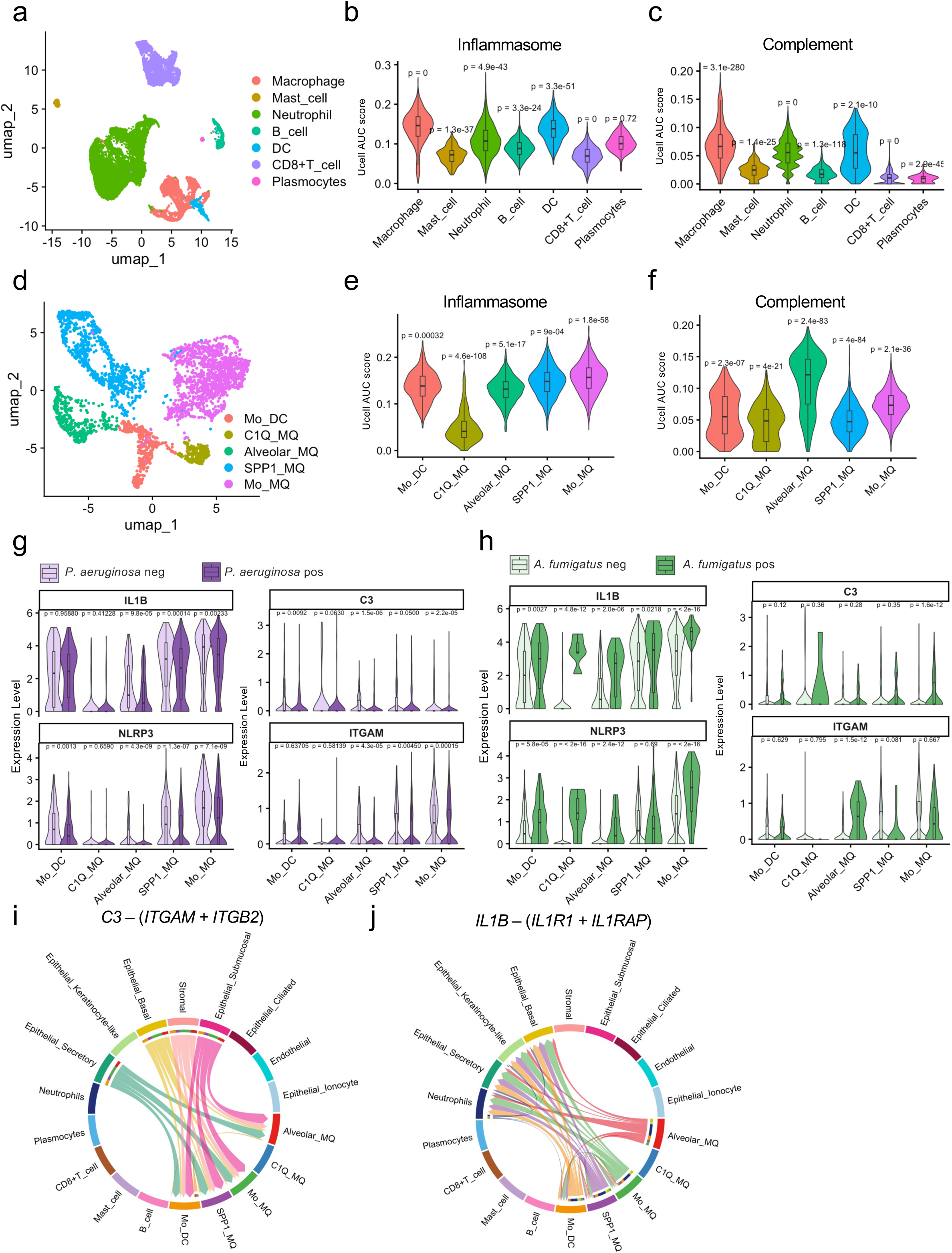
Single-cell transcriptomic profiling of inflammasome and complement expression in pulmonary cells from sputum and biopsy of people with cystic fibrosis (pwCF). **(a)** Uniform Manifold Approximation and Projection (UMAP) highlighting immune subclusters. **(b-c)** Violin plots comparing the expression scores of inflammasomes **(b)** and complement **(c)** pathways across major immune cell populations. **(d)** UMAP projection focusing on macrophages and dendric cell subset annotation (monocyte-derived dendritic cell (mo_DC), C1Q type macrophages (C1Q_MQ), alveolar macrophages (Alveolar_MQ), SPP1 type macrophages (SPP1_MQ) and monocyte-derived macrophage (Mo_MQ)). **(e-f)** Violin plots comparing the expression scores of inflammasomes **(e)** and complement **(f)** pathways across macrophages/DC cell populations. **(g-h)** Quantitative expression analysis comparing *NLRP3*, *IL1B*, *ITGAM* and *C3* levels between *P.a*-negative and -positive conditions **(g)** and *A.f*-negative and -positive conditions across macrophage and dendritic cell subsets **(h)**. **(i)** Chord diagram (CellChat analysis) showing inferred *C3*–CR3 (*ITGAM*–*ITGB2*) signaling among the annotated cell populations. **(j)** Chord diagram (CellChat analysis) showing inferred *IL1B*–(*IL1R1*/*IL1RAP*) signaling among the annotated cell populations.

Within the macrophage/DC compartment, we resolved distinct subsets based on canonical marker genes (Supplementary Fig. 8d), including monocyte-derived macrophages (Mo_MQ), C1Q macrophages (C1Q_MQ), alveolar macrophages (Alveolar_MQ), SPP1 macrophages (SPP1_MQ), and monocyte-derived DCs (Mo_DC) (Fig. 8d). Inflammasome pathway genes (Fig. 8e, Supplementary Fig. 8e), including *NLRP3* and *IL1B* were enriched in Mo_MQ and SPP1_MQ (Supplementary Fig. 8f). In contrast, complement pathway genes and our key components (e.g *C3* and *ITGAM*) were most prominent in Alveolar_MQ and Mo_MQ (Fig. 8f and Supplementary Fig. 8f-g). Across macrophage/DC subsets, *IL1B* expression correlated with *NLRP3* and *ITGAM* (Supplementary Fig. 8h), suggesting a strong functional link between the complement and inflammasome systems in these subsets.

To assess how microbial status shapes these pathways, we performed differential expression and pathway enrichment analyses stratified by infection group. GSEA indicated increased enrichment of inflammasome signaling and core-component gene sets in Mo_MQ from *P. aeruginosa*-positive pwCF, with reduced enrichment in Alveolar_MQ, SPP1_MQ, and Mo_DC (Supplementary Fig. 9a-e). Consistent with this pattern, *NLRP3* and *IL1B* were decreased in the same subsets (Fig. 8g). For the complement pathway, GSEA showed enrichment in Alveolar_MQ and C1Q_MQ, with reduced enrichment in SPP1_MQ in *P. aeruginosa*–positive pwCF, concordant changes were observed for *C3* and *ITGAM* (Fig. 8g).

Because *P. aeruginosa* positivity may reflect chronic colonization rather than acute infection, we next restricted analysis to exacerbation samples and compared *P. aeruginosa*-positive versus -negative pwCF. Interestingly in this context, inflammasome signaling and core component sets remained significantly enriched in *P. aeruginosa*-positive pwCF (Supplementary Fig. 9f).

Regarding fungal infection, pwCF co-positive for *A. fumigatus* and *Stenotrophomonas maltophilia*, another Gram-negative bacterium like *P. aeruginosa* (Supplementary Table 2), complement gene sets were broadly upregulated across macrophage and DC subsets compared with *A. fumigatus*–negative but bacterially positive individuals, independent of exacerbation status (Supplementary Fig. 10). Interestingly the C1Q_MQ subset showed concomitant enrichment of complement and inflammasome pathways in *A. fumigatus*–positive pwCF (Supplementary Fig. 10c). At the gene level, *IL1B* was increased across macrophage/DC subsets, whereas *NLRP3* increased in most subsets but not in SPP1_MQ. *C3* upregulation was most evident in Mo_DC, whereas *ITGAM* was most increased in Alveolar_MQ (Fig. 8h).

These patterns suggested that complement ligands and inflammasome activation may be distributed across distinct cellular sources and responders. To infer intercellular communication linking these programs, we performed CellChat analysis across the whole annotated cell populations. C3 signaling was predicted to originate predominantly from epithelial cells (secretory, basal, submucosal, and keratinocyte-like subsets) and stromal cells and to engage macrophage/DC populations primarily via the CR3 receptor complex (*ITGAM*–*ITGB2*) (Fig. 8i and Supplementary Fig. 11a,b). In contrast, *IL1B* signaling was predicted to arise mainly from Mo_MQ, Alveolar_MQ, SPP1_MQ, and Mo_DC and to target mainly neutrophils as well as epithelial subsets (secretory, basal, and keratinocyte-like) (Fig. 8j and Supplementary Fig. 11c,d).

Collectively, these results indicate that bacterial–*A. fumigatus* superinfection amplifies inflammatory gene signatures in pwCF and suggests coordinated engagement of complement- and inflammasome-related pathways across airway myeloid compartments. Notably, our analyses predicted epithelial/stromal to myeloid C3 signaling and, in turn, IL1B signaling from myeloid toward neutrophils and epithelial cells, consistent with a feed-back loop linking complement activation to IL-1–driven inflammation in CF airways and potentially contributing to dysregulated airway inflammation.

## DISCUSSION

Chronic polymicrobial infection is a defining feature of CF airways and a major driver of progressive lung inflammation and tissue damage^8^. Here, we uncover a mechanistic basis for the exaggerated inflammatory response that emerges during sequential infection with *P. aeruginosa* and *A. fumigatus*, two pathogens frequently co-colonizing the lung of pwCF^9–12^. We demonstrate that prior exposure to *P. aeruginosa* functionally potentiates macrophages to prepare an amplified inflammasome response upon subsequent fungal infection. Although increased IL-1β production has been observed during co-infection of *A. fumigatus* with *P. aeruginosa*^12^ or *Mycobacterium tuberculosis*^13^, the molecular mechanisms driving this exaggerated inflammatory response have remained undefined.

Strikingly, the amplified inflammasome response persisted even after *P. aeruginosa* was fully cleared prior to fungal exposure, indicating that sustained bacterial presence is not required for hyperactivation. This finding is particularly relevant given the uncertain spatial proximity of these pathogens within CF airways, but our results suggest that temporal separation does not prevent cross-kingdom inflammatory amplification. Importantly, this potentiation required active infection with live bacteria and could not be reproduced by stimulation with isolated *P. aeruginosa* PAMPs, demonstrating that the phenomenon extends beyond classical macrophage priming. Consistent with this interpretation, LPS exposure, but not flagellin, enhanced fungal control without promoting inflammasome overactivation. These findings support a model in which distinct bacterial signals establish multiple layers of macrophage conditioning, a likely TRIF-dependent pathway driven by TLR4 that improves controlling fungal growth, and a separate mechanism linked to live infection that licenses excessive inflammasome activation during subsequent *A. fumigatus* challenge.

This inflammasome hyperactivation during *P. aeruginosa*-*A. fumigatus* superinfection is independent of NLRC4 or AIM2 and instead proceeds through the canonical NLRP3–ASC–caspase-1/8 axis. Interestingly, IL-1β release occurred independently of gasdermin-D, consistent with recent observations in other infection contexts^14^. This finding suggests that inflammasome-driven cytokine secretion during bacterial–fungal superinfection may involve alternative, GSDMD-independent pathways. One possibility is a role for gasdermin-E, which can be activated by caspase-3 and has been implicated in non-canonical inflammatory responses, reinforcing this hypothesis, *A. fumigatus* has been shown to induce caspase-3 activation in BMDMs^15^. In addition, a potential mechanical stress or membrane disruption associated with fungal hyphal growth could contribute to cytokine release without requiring classical pore-forming proteins^16^. The precise mechanism enabling IL-1β secretion in the absence of gasdermin-D remains unclear and warrants further investigation, particularly given its potential relevance to chronic airway inflammation in pwCF.

Our findings reveal that this bacterial-fungal amplification of inflammasome activity depends on specific PAMPs from both pathogens. On the bacterial side, flagellin, the type III secretion system, and type IV pili proved essential for macrophage potentiation. These virulence factors are known inflammasome activators during *P. aeruginosa* infection^17–20^, supporting their continued importance in driving inflammasome hyperactivation even after bacterial clearance. On the fungal side, the cell-wall polysaccharide GAG emerged as a critical determinant of inflammasome activation, consistent with our previous observations during *A. fumigatus* infection^2^. Together, these microbial components establish a signaling landscape that selectively amplifies IL-1β production. Importantly, this phenotype was reproduced using matched microbial clinical CF isolates, underscoring the relevance of this mechanism within the complex microbial environment of the airway of the pwCF.

Mechanistically, our data identify the MyD88 signaling adaptor as a central regulator of this hyperinflammatory response. This result confirms our previous observation highlighting a response to pathogens engaging a multilayer of signaling cascade necessary for an efficient immune response between controlling the fungal growth and engaging inflammasome pathway. MyD88 deficiency in murine or human macrophages abrogated caspase-1 activation and IL-1β release, implicating TLR-dependent rather than IL-1R–mediated signaling in inflammasome regulation during bacterial-fungal superinfection. Transcriptomic analyses of *Myd88^−/−^* macrophages revealed a distinct loss of infection-induced transcriptional reprogramming, identifying ITGAM as a potential downstream effector.

Functional experiments demonstrated that ITGAM is essential for inflammasome hyperactivation during bacterial-fungal superinfection. While ITGAM has previously been reported to negatively regulate NLRP3 activation in response to monosodium urate crystals, likely through a metabolic shift^21^, its role appears to be highly context dependent. In ischemic stroke, where NLRP3-driven inflammation exacerbates tissue damage^22,23^, ITGAM instead promotes IL-1β production^24^, consistent with our findings in macrophages exposed to sequential *P. aeruginosa* and *A. fumigatus* infection. Notably, matrix metalloproteinase-9 (MMP9), highlighted in that setting as an ITGAM-associated inflammatory factor^24^, was similarly upregulated in our transcriptomic dataset, suggesting convergence on shared pathogenic signaling pathways across these distinct inflammatory conditions.

ITGAM is a key innate immune integrin, and our data indicate that its relevance in bacterial-fungal superinfection derives from its role as a complement receptor. In sequentially infected macrophages, complement pathway genes were strongly upregulated, and functional studies demonstrated that ITGAM cooperates with its binding partner ITGB2 to engage C3. Consistently, BMDMs lacking C3 showed markedly impaired caspase-1 activation and IL-1β release, establishing that ITGAM and its ligand C3 constitute a critical signaling axis that links complement activation to inflammasome hyperactivation in this context. Concomitantly, blocking ITGAM or deleting C3 phenocopied the defective inflammasome activation seen in *Myd88^−/−^* cells, accompanied by similar reduction of SYK phosphorylation. These findings establish a previously unrecognized MyD88–ITGAM–C3–SYK signaling cascade that coordinates innate immune amplification during sequential bacterial-fungal challenge.

Consistent with our findings, mice lacking C3 exhibit increased susceptibility to *P. aeruginosa* pneumonia^25^ and *A. fumigatus* infection^26^, underscoring the importance of complement in host defense against these two pathogens. Moreover, complement-mediated opsonization of *A. fumigatus* conidia, particularly via C3b, was recently shown to enhance IL-1β production in human monocyte-derived macrophages^27^, suggesting a broader role for complement in inflammasome regulation during fungal infection. Our study further highlights the contribution of fungal GAG to inflammasome activation in the setting of bacterial-fungal co-infection. Future work will be required to delineate how GAG exposure integrates with complement receptor engagement to coordinate C3-dependent signaling and downstream inflammasome amplification.

Our scRNA-seq analysis of sputum and lung biopsy samples from pwCF, stratified by *P. aeruginosa* and *A. fumigatus* status, provides translational support for this pathway in human disease. Inflammasome- and complement-associated genes were predominantly expressed in macrophages, dendritic cells (DCs), and neutrophils. Pathway enrichment analyses further suggested that co-positivity for *A. fumigatus* and Gram-negative bacteria is associated with a coordinated inflammatory transcriptional program in C1Q_MQ subset, characterized by concomitant upregulation of complement and inflammasome pathways. These findings mirror our *in vitro* results and support the clinical relevance of complement-dependent potentiation of inflammasome responses in CF airways. However, the cellular sources of complement signals and inflammasome-associated responses appear to be at least partly distinct in pwCF airways. Indeed, CellChat analysis predicted a putative epithelial/stromal-to-myeloid signaling axis, in which epithelial and stromal cells act as major sources of C3 signaling that may engage CR3 on macrophages and DCs, which in turn are predicted to generate IL-1 signaling toward neutrophils and epithelial cells. This feed-back loop provides a plausible framework for persistent and dysregulated airway inflammation in airway of pwCF.

Taken together, our findings support a model in which sequential infection with *P. aeruginosa* or Gram-negative bacteria and *A. fumigatus* drives a pathogenic amplification of macrophage inflammasome responses in airways. During the initial bacterial encounter, TLR engagement activates MyD88- and TRIF-dependent signaling pathways, promoting macrophage activation and upregulation of ITGAM as part of a protective program that contributes to inflammatory resolution post bacterial infection^28^. However, within the polymicrobial environment of the CF lung, subsequent exposure to *A. fumigatus* leads to complement activation and C3 deposition on fungal surfaces. Recognition of these C3 fragments by the ITGAM–ITGB2 (CR3) receptor complex reinforces SYK-dependent intracellular signaling, thereby licensing excessive activation of the canonical NLRP3 inflammasome and driving disproportionate IL-1β release into the lung. Such inflammatory amplification likely contributes to the persistent, tissue-damaging immunopathology observed in pwCF experiencing bacterial–fungal co-infection.

Collectively, this work defines a mechanistic framework in which bacterial infection reshapes macrophage signaling and transcriptional responses, potentialize them to subsequent fungal stimuli. The identification of the ITGAM–C3 axis as a key driver of NLRP3 inflammasome hyperactivation reveals an unexpected convergence between complement and inflammasome pathways in the context of bacterial and fungal superinfection.

From a translational perspective, these findings suggest that targeting complement signaling or specifically the ITGAM–C3 interaction could attenuate pathological inflammasome activation without compromising essential host defense. Future work should explore whether modulation of this axis can restore immune balance and mitigate inflammation-driven lung damage in pwCF.

## EXPERIMENTAL PROCEDURES

### Mice

Wild-type C57BL/6 mice were purchased from R. Janvier (Le Genest Saint-Isle, France). Transgenic mouse strains including *Myd88^−/−^*, *Ifnar1^−/−^*, *Tnfa^−/−^*, *Nlrp3^−/−^*, *Aim2^−/−^* and *Gsdmd^−/−^* were bread at the animal facility: Typage et Archivage d’Animaux Modèles (TAAM) UAR44, CNRS, Orléans, France. All mice were backcrossed to the C57BL/6 background. *C3^−/−^* mice was bred provided by Complement Therapeutics Research Group and National Renal Complement Therapeutics Centre (Newcastle University Translational and Clinical Research Institute, Newcastle, United Kingdom).

### Microbial Cultures

***Pseudomonas aeruginosa.*** Different strains of *Pseudomonas aeruginosa* (Table 1 **| Pseudomonas aeruginosa strains used in this study** were cultured on Lysogeny Broth (LB) agar plates and incubated overnight at 37°C. The day before infection, a single colony was selected and inoculated into 5 mL of liquid LB medium for 8 hours. Serial dilutions (10-fold) were performed up to 10⁻¹¹ and incubated for 16 hours at 37°C to reach an optical density at 600 nm (OD600) between 0.3 and 0.7, corresponding to the exponential bacterial growth phase. Macrophages were infected at a multiplicity of infection (MOI) of 0.05, 0.01, or 0.005.

**Table 1.** *Pseudomonas aeruginosa* strains used in this study.

| Bacterial Strain | Wild-Type Background | Phenotype | Reference |
| --- | --- | --- | --- |
| PAO1 | Clinical isolate<br>CECT<br>4122/ATCC<br>15,692 | – |  |
| PAK | Wild-type strain | – |  |
| $\Delta pscF$ | PAK | Mutant for the Type III Secretion System (T3SS), no longer secretes exotoxins S, T, Y, and U | 29 |
| $\Delta flic$ | PAK | Mutant for flagellin | 30 |
| $\Delta flic\Delta pscF$ | PAK | Double mutant for flagellin and Type III Secretion System (T3SS) | 29 |

***Aspergillus fumigatus.*** Conidia from different *Aspergillus fumigatus* strains (Table 2 **| Aspergillus fumigatus strains used in this study** were cultured on inclined Sabouraud 4% Glucose Agar and incubated at 35°C for 6 days. Prior to infection, conidia were harvested in 13 mL of phosphate-buffered saline (PBS) containing 0.1% Tween-20, then filtered twice using a 40-μm cell strainer. Macrophages were infected at an MOI of 5 (five conidia per macrophage).

**Table 2.**
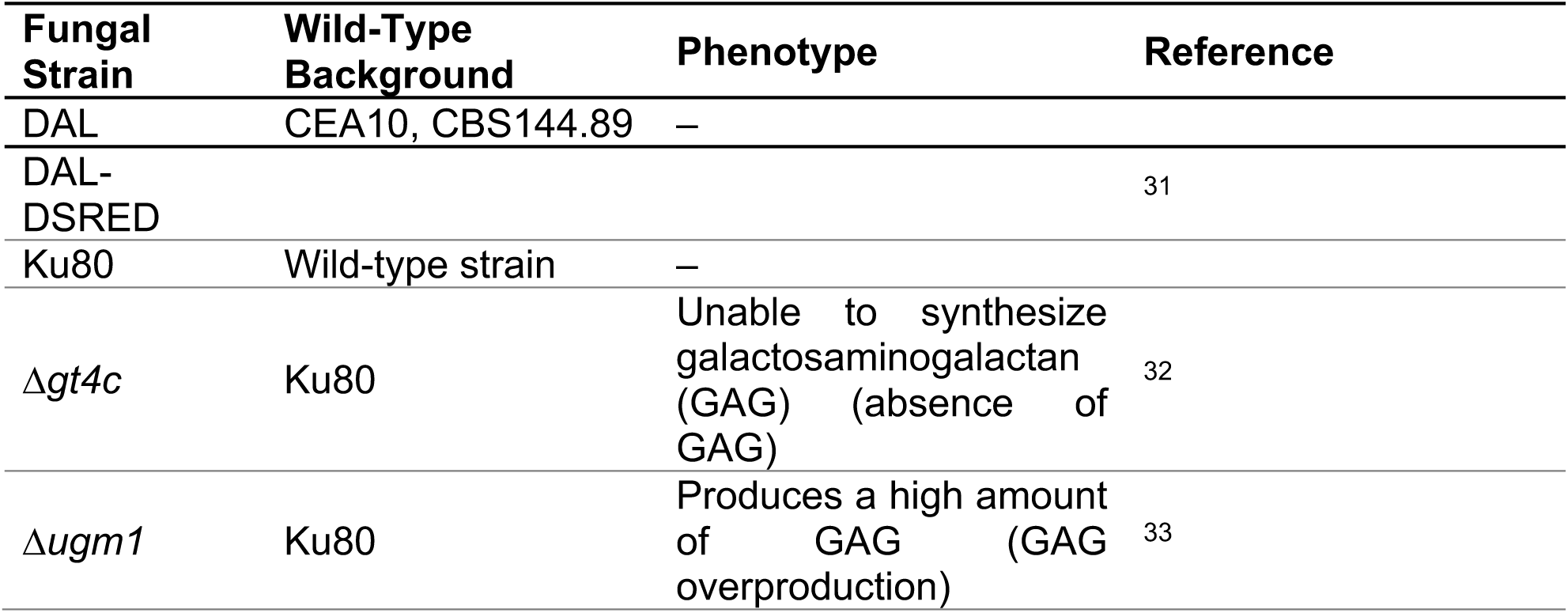
*Aspergillus fumigatus* strains used in this study.

**Table 3.** List of flow cytometry antibodies used in this study.

| Anti-mouse antibodies | Fluorophore | Clone | Source | # | Dilution |
| --- | --- | --- | --- | --- | --- |
| CD45 | BV605 | 30-F11 | BIOLEGEND | 103155 | 1/400 |
| Ly6G | FITC | 1A8 | BIOLEGEND | 127606 | 1/200 |
| CD11c | Pe-Cy7 | N418 | BIOLEGEND | 117318 | 1/800 |
| F4/80 | Pe | BM8 | BIOLEGEND | 123110 | 1/200 |
| I-A/I-E | PerCP | M5/114,15,2 | BIOLEGEND | 107623 | 1/400 |
| CD64 | APC | X54-5/7.1 | BIOLEGEND | 139306 | 1/200 |
| CD11b | APCvio7 | M1/70 | BIOLEGEND | 101226 | 1/800 |

### Cell Cultures. Bone Marrow-Derived Macrophages (BMDMs)

Mice were euthanized, and femurs and tibias were harvested. Bone marrow was flushed using sterile PBS and red blood cells lysed by adding 10 mL of 0.2% NaCl for 30 seconds, followed by osmotic balance restoration with 10 mL of 1.6% NaCl. Cells were centrifuged at 400g for 5 minutes. The remaining cells were resuspended and cultured in 20-cm Petri dishes (639160, Greiner) in macrophage differentiation medium consisting of RPMI 1640 with GlutaMAX supplement (61870-010, Gibco), 15% 3T3-conditioned medium (containing macrophage colony-stimulating factor [M-CSF]), 10% fetal bovine serum (FBS), 1% non-essential amino acids (NEAA), 1% penicillin-streptomycin (P/S) and 1x β-mercaptoethanol (21985023, Sigma). Cells were cultured at 37°C in 5% CO_2_ for 7 days, with 5 mL of differentiation medium added on days 3 and 5. On day 6, cells were seeded at 1×10⁶ macrophages per well in 12-well plates (Falcon) for infection on day 7.

### Murine Alveolar Macrophages (mAMs)

Murine alveolar macrophages were isolated from C57BL/6J mice by performing two bronchoalveolar lavages with 1 mL of PBS at 37°C. The collected lavage fluid was centrifuged at 300g for 5 minutes, and the pellet was resuspended in culture medium (RPMI, 10% FBS, 1% P/S, 1% NEAA, and 10 ng/mL GM-CSF (#315-03, PEPROTECH). Cells were maintained at 37°C in 5% CO_2_ for 7 days before being seeded for infection experiments as described for BMDMs.

### *In vitro* stimulation

On day 7, cells were washed with PBS and incubated in DMEM with 10% FBS without antibiotics. After 2 hours of resting, macrophages were infected with *P. aeruginosa* at the previously described MOI. After 4 hours of bacterial infection, cells were washed with PBS and incubated in DMEM (10% FBS) supplemented with 1% P/S and 80 μg/mL Amikacin (A2324-5G, Sigma). The washing step was repeated 2 hours later. Cells were incubated for 22 hours at 37°C in 5% CO_2_. After 22 hours, macrophages were washed with PBS and infected with *A. fumigatus* at an MOI of 5 in the presence of antibiotics for 16 hours at 37°C in 5% CO2. For priming experiments, BMDMs were incubated with 100 ng/mL LPS (PA10 L9143, Sigma-Aldrich), 1 μg/mL flagellin (tlrl-pafla, Invivogen), or 100 ng/mL IFN-γ (P01579, Invivogen) for 4 hours before *A. fumigatus* infection.

### Heat killed / PFA fixed *Pseudomonas aeruginosa*

*Pseudomonas aeruginosa* PAO1 was cultured and harvested at the desired growth phase, as described in the previous Microbial Cultures section. For PFA fixation, bacterial cultures were fixed overnight with 4% PFA at 4°C. The fixed bacteria were then washed three times in PBS containing 0.1M NH_4_Cl to quench residual PFA, followed by two additional washes with PBS to remove any remaining reagent. For heat inactivation, harvested bacteria cells were washed three times in PBS and subsequently incubated at 60°C for 40 minutes. To verify successful inactivation, both PFA-fixed and heat-killed bacteria were plated on LB agar and incubated at 37°C for 24-48 hours to confirm the absence of bacterial growth.

### Phagocytosis and Conidiocidal Assays

Phagocytosing ability per BMDM was determined by calculating the number of BMDMs that had one or more conidia phagocytosed after 1 hour of infection with 10 MOI of *A. fumigatus* resting conidia. Following this, cells were lysed with 0.1 % triton X100, plated on LB petri dishes and the number of colonies forming units was counted.

### Bacterial CFU

To quantify intracellular bacterial burden, bone marrow-derived macrophages (BMDMs) were infected with *Pseudomonas aeruginosa* at a multiplicity of infection (MOI) of 0.05, 0.01, or 0.005. After 1 hour of infection at 37°C and 5% CO₂, extracellular bacteria were removed by washing the cells twice with sterile phosphate-buffered saline (PBS), followed by incubation with culture medium containing 50 µg/mL amikacin for 1 hour to kill remaining extracellular bacteria. Subsequently, macrophages were lysed using 0.1% Triton X-100 in PBS for 5 minutes at room temperature. The lysates were serially diluted in PBS and plated on LB agar plates. Plates were incubated overnight at 37°C, and colony-forming units (CFUs) were counted to determine the number of viable intracellular bacteria.

### Inhibitor Treatments and blocking antibodies

To evaluate inflammasome activation, BMDMs were treated with various inhibitors at the time of *A. fumigatus* superinfection, NLRPC inhibitor MCC950 (10 μM final concentration) (inhibit-mcc, InvivoGen), caspase-1 and caspase-4 inhibitor VX-765 (40 μM final concentration) (inh-vx765i-1, InvivoGen), caspase-8 inhibitor Z-IETD-FMK (20 μM final concentration) (inhibit-ietd, InvivoGen). To evaluate CD11b implication, BMDMs were pretreated with either 20µg/mL of inhibitory monoclonal antibody against CD11b (clone M1/70) (#14-0112-86; eBioscience, Invitrogen) or 20µg/mL of IgG2b control (#15277347, eBioscience™, Invitrogen™) at 37°C and 5% CO_2,_ 3 h prior to *A. fumigatus* infection and maintained during the rest of the infection.

### ELISA Assay

Cytokine secretion was analyzed by ELISA for mouse IL-1β (DY401, R&D Systems), TNF-α (DY410, R&D Systems), and KC (DY453, R&D Systems). All ELISA were performed according to the manufacturer’s instructions. Absorbance at 450 nm was measured using a Multiskan Sky microplate spectrophotometer (Thermo Fisher Scientific) at the final step.

### Western Blot Analysis

For the detection of caspase-1 cleavage, both cell pellets and culture supernatants were lysed in caspase lysis buffer consisting of 1 tablet of protease inhibitor cocktail (Roche, 11697498001), 200 µL of 1 M dithiothreitol (DTT; Roche, 10197777001), 2 mL of NP-40 (Millipore, 492016), and Milli-Q water. For the analysis of signaling pathway molecules, BMDMs were lysed in 1× RIPA buffer supplemented with one PhosSTOP phosphatase inhibitor tablet (Roche, 04906837001) and one protease inhibitor tablet (cOmplete™, Roche, 04963116001) per 10 mL of buffer. Protein lysates were denatured by boiling at 95°C for 10 minutes and separated on 13% SDS-PAGE gels. Proteins were transferred to 0.2 µm nitrocellulose membranes (Amersham™, 10600006), followed by blocking in TBS-Tween 0.01% containing either 5% skimmed milk or 3% bovine serum albumin (BSA), depending on antibody requirements. Primary antibodies were incubated overnight at 4°C at a 1:1000 dilution, followed by incubation with horseradish peroxidase-conjugated secondary antibodies (anti-mouse, Jackson ImmunoResearch, 315-035-047; or anti-rabbit, Jackson ImmunoResearch, 111-035-047) at a 1:5000 dilution. Antibodies used include anti-caspase-1 (AG-20B-0042-C100, ADIPOGEN), anti-pIκBα (2859S, Cell Signaling Technologies [CST]), anti-tIκBα (4814S, Cell Signaling Technologies [CST]), anti-pERK (9101S, Cell Signaling Technologies [CST]), anti-tERK (9102S, Cell Signaling Technologies [CST]), anti-pSYK (pZap70) (65E4, Cell Signaling Technologies [CST]), anti-tSYK (2712S, Cell Signaling Technologies [CST]), anti-pro-IL1β (D3H12, Cell Signaling Technologies [CST]), anti-NLRP3 (D4D8T, Cell Signaling Technologies [CST]), and anti-β-actin (A3854, Sigma Aldrich). Membranes were developed by chemiluminescent detection using the ChemiDoc Imaging System (Bio-Rad), with ECL Clarity substrate for HRP (Biorad, #1705060) and analyzed with ImageLab software (Bio-Rad).

### Real-Time Cell Death and Fungal Growth Analysis (live cell imaging analysis)

Real-time monitoring of cell death and fungal growth was performed using the IncuCyte® live-cell imaging system (IncuCyte SX5, Sartorius™, Göttingen, Germany) maintained at 37°C with 5%vfds CO₂. Cell nuclei of dying cells were labeled using propidium iodide (PI) (cat.no. P4170-25MG, Sigma-Aldrich) at a final concentration of 1 µMor with SytoxGreen (S7020, Invitrogen) diluted in 1:2500 into the culture medium throughout the experiment.

Time-lapse images were acquired automatically at regular intervals and analyzed using the IncuCyte SX5 integrated analysis software. Quantification of fungal hyphal growth (*Aspergillus* DAL-DSRED strain, Table 2), was performed using the *NeuroTrack* analysis module of the IncuCyte SX5 software^34,35^.

### RNA-Sequencing analysis

Quality Control (QC) of raw sequencing data was performed using FastqQC FastQ (v0.11.8; https://github.com/s-andrews/FastQC) and MultiQC (v1.31,^36^). Reads alignment and transcript quantification were performed using Kallisto (v0.51.0)^37^. Differential Expression Analysis (DEA) was carried out on raw count data using DESeq2 R package^38^. Principal Component Analysis (PCA) were generated from the normalized data from DESeq2. Results were visualized using a Heatmap from the heatmap (https://github.com/raivokolde/pheatmap) R package. Finally, a Gene Ontology (GO) enrichment analysis was performed using the ShinyGO 0.85.1^39^.

### Single cell RNA-sequencing analysis

We reprocessed single-cell RNA-seq data from twelve samples obtained from pwCF, including three lung explants^40^ and nine sputum specimens (GSE145360)^41^ (Supplementary Table 2). Raw counts, h5ad object and metadata were downloaded from GEO40 and Cellxgene39 databases. Seurat objects were created using Seurat v5.3.1, samples were filtered to only contain cells with 200 or more expressed genes, <15 % mitochondrial RNA. Samples from the two datasets were merged and 3000 highly variable genes identified using the “vst” method. Thereafter datasets were integrated using the RCPA method (k.anchor=15). Gene signature scores were calculated using the AddModuleScore_UCell fonction. CellChat v2.2.0 was used to infer cell-cell interactions. For further details regarding the materials and methods used, please refer to the Supplementary Materials and Methods.

### STRING analysis

The string analysis was performed by using the version 12.0 available on https://string-db.org/^42^.

### Flow Cytometry

Cells were resuspended in FACS buffer (phosphate-buffered saline [PBS] supplemented with 2 mM EDTA and 2% fetal bovine serum [FBS]) and incubated with anti-mouse CD16/CD32 Fc-blocking antibody (clone 2.4G2; #553142, BD Biosciences) for 20–30 minutes at 4°C to prevent nonspecific binding. Following Fc receptor blockade, cells were washed with FACS buffer and stained with the LIVE/DEAD™ Fixable Blue viability dye (#L23105, Thermo Fisher Scientific; 1:1000 dilution) and a panel of fluorochrome-conjugated monoclonal antibodies for 30 minutes on ice in the dark. Antibodies used include CD45 (clone 30-F11, BV605; BioLegend #103155, 1:400 dilution), Ly6G (clone 1A8, FITC; BioLegend #127606, 1:200 dilution), CD11c (clone N418, PE-Cy7; BioLegend #117318, 1:800 dilution), F4/80 (Pe; clone BM8, PE; BioLegend #123110, 1:200 dilution), I-A/I-E (MHC II) (clone M5/114.15.2, PerCP; BioLegend #107623, 1:400 dilution), CD64 (FcγRI) (clone X54-5/7.1, APC; BioLegend #139306, 1:200 dilution), and CD11b (clone M1/70, APC-Cy7; BioLegend #101226, 1:800 dilution). After staining, cells were rewashed with FACS buffer. Data acquisition was performed on a CytoFLEX flow cytometer (Beckman Coulter), and analyses were conducted using FlowJo software (version 10.4.2, BD Biosciences).

### 13. Statistical analysis

All statistical analyses were conducted using GraphPad Prism software (version 10.4.1, GraphPad Software, San Diego, CA, USA). Data are presented as indicated in the respective figure legends, including the statistical tests applied. The number of biological replicates (n) used for each experimental condition is also specified in the figure legends. Significance levels and error bars are defined accordingly.

## Acknowledgements

This work was supported by the patient associations “Vaincre la Mucoviscidose” (VLM) and “Association Gregory Lemarchal”: RF20230503265 (to BB and LG, NR), RF20240503504 (to BB). BB was also supported by the “Agence National de la Recherche” (ANR): ANR-23-CE15-0013-03, ANR-24-CE15-2995-03 and ANR-25-CE15-1868-01. The authors thank Vishukumar Aimanianda and Nathalie Winter for helpful discussions throughout this study as CSI members for SK thesis. The authors also thank Valérie Quesniaux, Bernhard Ryffel and the TAAM UPS44 (Orléans, France) for their help with the mice.

---

**Supplementary Fig. 1| (a)** Immunoblot analysis showing caspase-1 processing (pro–CASP1 and p20) and **(b)** ELISA quantification of IL-1β secretion in BMDMs following infection with two wild-type *P.a* strains (MOI 0.005 or 0.01), PAO1 and PAK, under single or superinfection conditions with *A.f* (MOI 5). **(c, d)** Quantification of *A.f* hyphal growth (fungal length, mm/mm²) over time following macrophage priming with either PAO1 **(c)** or PAK **(d)** (at MOI 0.01), showing significant inhibition of fungal extension after bacterial pre-infection compared to *A.f* alone. **(e)** Enumeration of *P.a* CFUs at two multiplicities of infection (MOI 0.005 and 0.01) for WT and indicated mutant strains, assessed 1 hour post-infection. **(f)** Immunoblot analysis showing caspase-1 processing (pro–CASP1 and p20) in BMDMs following infection with *P.a* (MOI 0.05), *Escherichia coli* (*E. coli*) or *Staphylococcus aureus* (*S. aureus*) (MOI 0.1), under single or superinfection conditions with *A. fumigatus* (*A.f*) (MOI 5).

**Supplementary Fig. 2| NLRP3 hyperactivation involves caspase-1 & -8. (a)** Time-course quantification of cell death PI+ BMDM nuclei in WT and *Nlrp3*^−/−^ BMDMs left uninfected (Med) or after infection with *P.a* (MOI 0.01), and/or *A.f* (MOI 5). **(b)** Immunoblot analysis of caspase-1 processing (pro–CASP1 and p20) in BMDMs treated with or without the NLRP3 inhibitor MCC950 under the indicated infection settings. ELISA quantification of IL-1β **(c)** and TNFα **(d)** secretion by BMDMs ± MCC950. **(e)** Time-course quantification of cell death PI+ BMDM nuclei in WT and *Casp1*^−/−^ BMDMs left uninfected (Med) or after infection with *P.a* (MOI 0.01), and/or *A.f* (MOI 5). **(*f*)** Immunoblot analysis showing caspase-1 processing (pro–CASP1 and p20) in BMDMs from WT, *Casp1^−/−^*mice left uninfected (Med) or after infection with *P.a* (MOI 0.005 or 0.01), and/or *A.f* (MOI 5) and **(*g*)** corresponding IL-1β quantification for *Casp1^−/−^***. (h)** Time-course of fungal hyphal growth (length/mm²) in WT and *Casp1^−/−^* BMDMs left uninfected (Med) or after infection with P.a (MOI 0.01), and/or A.f (MOI 5). **(i)** IL-1β measurement by ELISA in BMDMs treated with pan-caspase inhibitor ZVAD-FMK, left uninfected (Med) or after infection with *P.a* (MOI 0.005 and 0.01), and/or *A.f* (MOI 5). **(j)** Quantification of *A.f* hyphal growth (length/mm² over time) in WT and *Gsdmd*^−/−^ BMDMs under the same infection conditions. All data represent mean ± SEM from at least three independent experiments. Statistical significance: *p<0.05, **p<0.01.

**Supplementary Fig. 3| Principal component analysis (PCA) of transcriptomic profiles following infection. (a)** PCA plot displaying sample clustering based on all expressed genes, colored by timepoint post-infection (0 h, 28 h corresponding to the pre-fungal stage, 32 h and 36 h corresponding to 4 h and 8 h post-fungal infection). Each point represents an individual sample at the indicated time. **(b)** PCA plot of the same dataset, colored by infection condition (*P. aeruginosa* pre-exposure (P*.a*) or without it (no *P.a*)). Data represent biological triplicate per group.

**Supplementary Fig. 4| Inflammasome overactivation is not dependent on TNF nor IFNAR1 pathway. (a)** Immunoblot analysis showing caspase-1 processing (pro–CASP1 and p20) and ELISA quantification of IL-1β **(b)** and TNFα **(c)** secretion in BMDMs from WT, *Tnf^−/−^* mice left uninfected (Med) or after infection with *P.a* (MOI 0.005 or 0.01), and/or *A.f* (MOI 5). **(d)** Immunoblot analysis showing caspase-1 processing (pro–CASP1 and p20) and ELISA quantification of IL-1β **(e)** and TNFα **(f)** secretion in BMDMs from WT, *Ifnar1^−/−^* mice left uninfected (Med) or after infection with *P.a* (MOI 0.005 or 0.01), and/or *A.f* (MOI 5). Statistical significance assessed by ANOVA; *p<0.05, ns = not significant.

**Supplementary Fig. 5| Inflammasome overactivation is not dependent on IL-1R pathway. (a)** ELISA quantification of KC secretion in BMDMs from WT and *Myd88^−/−^* mice left uninfected (Med) or after infection with *P.a* (MOI 0.005 or 0.01), and/or *A.f* (MOI 5). Immunoblot analysis showing caspase-1 processing (pro–CASP1 and p20 cleavage) in WT THP1 Dual and *Myd88^−/−^* THP1 macrophages under the indicated infection conditions. GAPDH serves as the loading control. **(c)** Immunoblot analysis showing caspase-1 processing (pro–CASP1 and p20 cleavage) and ELISA quantification of IL-1β **(d)** and TNFα **(e)** in BMDMs from WT and *Il1r1^−/−^* mice left uninfected (Med) or after infection with *P.a* (MOI 0.005 or 0.01), and/or *A.f* (MOI 5). **(f)** Quantification of cell death, measured as the number of SytoxGreen positive (Sytox^+^) cells, in *Il1r1^−/−^* BMDMs following infection with *P.a* (MOI 0.01) and/or *A.f* (MOI 5). **(g)** Time-course analysis of fungal growth (hyphal length in mm/mm²) of *A.f* (MOI 5) after *Il1r1^−/−^* BMDMs infection, either left uninfected (medium control, Med) or co-infected with *P.a* (MOI 0.005 or 0.01).

**Supplementary Fig. 6| Transcriptomic and functional characterization of *Myd88^−/−^*macrophage responses to *P. aeruginosa*. (a)** PCA plot displaying sample clustering based on all expressed genes in *Myd88^−/−^* macrophages, colored by timepoint post-infection (0 h, 28 h, 32 h, 36 h). Each point represents an individual sample at the indicated time. **(b)** PCA plot of the same dataset, colored by infection condition (*P. aeruginosa* pre-exposure (*P.a*) or without it (no *P.a*)) in *Myd88^−/−^*macrophages. **(c)** Heatmap showing the top 30 up- and downregulated genes in *Myd88^−/−^* BMDMs across samples, stratified by condition (*P. aeruginosa* pre-exposure (*P.a*) vs no *P.a*) and time point (32 h = 4 h post–*A. fumigatus* infection; 36 h = 8 h post–*A. fumigatus* infection). **(d)** Protein-protein interaction (STRING) network of the forty WT-specific candidate genes related to immune and inflammatory responses. Network nodes indicate proteins encoded by the candidate genes, and edges represent known or predicted interactions between them. The thickness of each edge reflects the strength or confidence of the interaction, with thicker edges representing stronger associations.

**Supplementary Fig. 7| Transcriptomic analysis revealed strong upregulation of *C3* among the top differentially expressed genes**. Heatmap of complement and immune-related gene expression across conditions and timepoints. Gene expression profiles of complement pathway components and selected immune genes were clustered and visualized as a heatmap. Each row represents a single gene, and each column represents an individual sample from wild-type (WT) or *Myd88^−/−^* knockout (KO) groups at specified infection conditions (infected with *P.a* (PA) corresponding to 28 hour time point, or without (sansPA)) followed by secondary infection with *A. fumigatus* for 4 or 8 hours (corresponding to 32- and 36-hour time points) and timepoints (as indicated on the x-axis). Color scale indicates row Z-score of normalized expression levels, with blue corresponding to lower, and red to higher, relative expression.

**Supplementary Fig. 8| Inflammasome and complement pathway activity across lung immune cell populations in two independent scRNA-seq datasets. (a)** Uniform Manifold Approximation and Projection (UMAP) plot showing major cell populations identified in scRNA-seq data. **(b,c)** Violin plots of inflammasome **(b)** and complement **(c)** pathways across the major cell populations. **(d)** Dot plot showing expression of selected marker genes (*FCN*, *VCAN*, *IL1B*, *SPP1*, *ITGB8*, *MMP19*, *MRC1*, *APOE*, *FABP4*, *HLA-DPA1*, *CD74*, *C1QA*, *CD1C*, *FCER1A*, *CLEC10A*) across macrophage and dendritic cell subsets. Dot size indicates the fraction of cells expressing each gene and color denotes average expression. **(e-f)** UMAP plots showing spatial distribution and expression intensity of inflammasome **(e)** and complement **(f)** gene expression signatures across all macrophages and DCs. **(g)** UMAP plots showing spatial distribution and expression intensity of key genes in inflammasome component (*NLRP3, IL1B*) and complement pathways (*ITGAM*, *C3*), across macrophage and dendritic cell subsets. **(h)** Correlation of expression levels between *IL1B* and *ITGAM*_*NLRP3* shown for macrophage and dendritic cell subclusters (scatter plot with regression line).

**Supplementary Fig. 9| Pathway enrichment analysis in single-cell macrophage and DC subsets from pwCF airways with *P. aeruginosa* (in stable or exacerbate status). (a-f)** Bar plots showing normalized enrichment scores for hallmark pathways in **(a)** Mo_MQ subcluster comparing *P.a* positive *vs.* negative subjects, and in **(b)** Alveolar_MQ, C1Q_MQ **(d)** SPP1_MQ, and **(e)** Mo_DC subclusters. **(f)** Bar plots showing normalized enrichment scores for hallmark pathways comparing *P.a* positive but in exacerbate status versus negative subjects. Enrichment directions are shown in blue for negatively enriched (downregulated) pathways and in red for positively enriched (upregulated) pathways.

**Supplementary Fig. 10| Pathway enrichment analysis in single-cell macrophage and DC subsets from pwCF airways with *Aspergillus*-positive samples.** Bar plot showing normalized enrichment scores in **(a)** Mo_MQ, **(b)** Alveolar_MQ, **(c)**, C1Q_MQ SPP1_MQ and **(e)** Mo_DC subclusters for hallmark pathways significantly upregulated (red bars) in *Aspergillus*-positive samples. Pathways are ordered by enrichment score on the x-axis.

**Supplementary Fig. 11| CellChat analysis reveal dialog between epithelial and myeloid compartments for inflammasome activation by complement. (a,b)** Heat map showing the **(a)** communication probability and **(b)** importance of the complement signaling network in each cell subtype. **(c,d)** Heat map showing the **(c)** communication probability and **(d)** importance of the IL-1 signaling network in each cell subtype.

