## Supplementary Figures for "Hijacking of inflammasome responses by the complement system during *Pseudomonas aeruginosa*–*Aspergillus fumigatus* sur-infection"

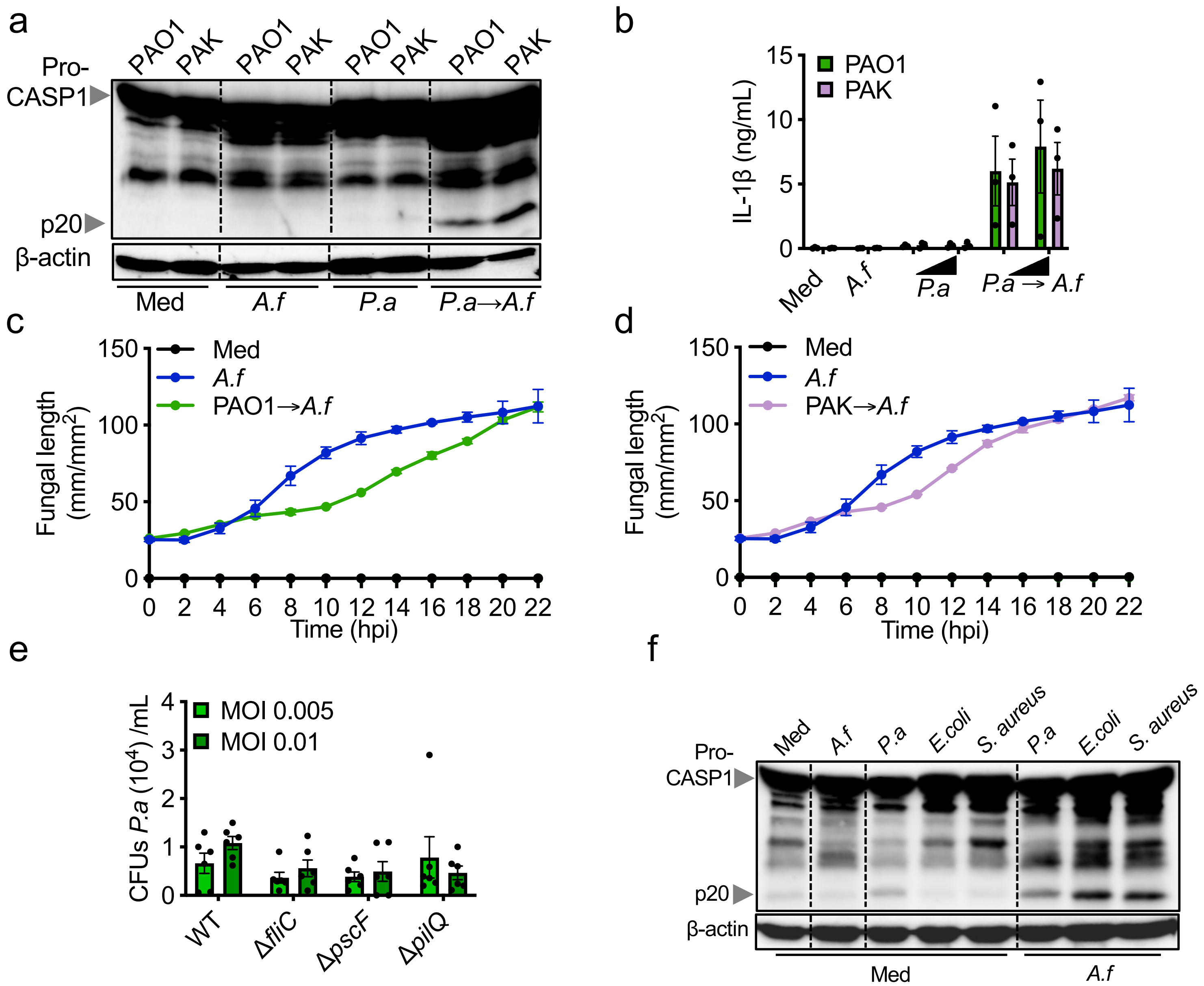


**Supplementary Fig. 1 |**

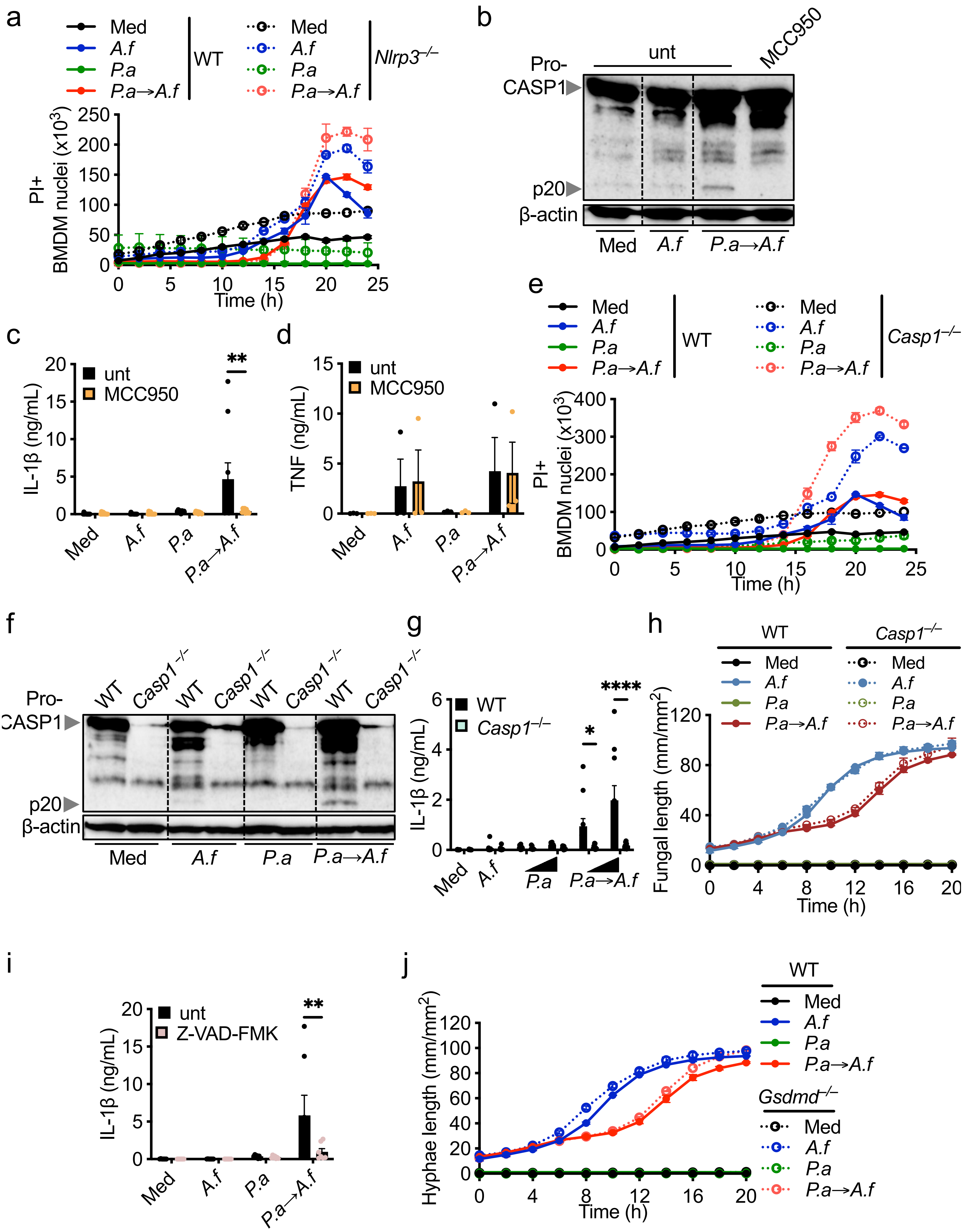


**Supplementary Fig. 2 |**

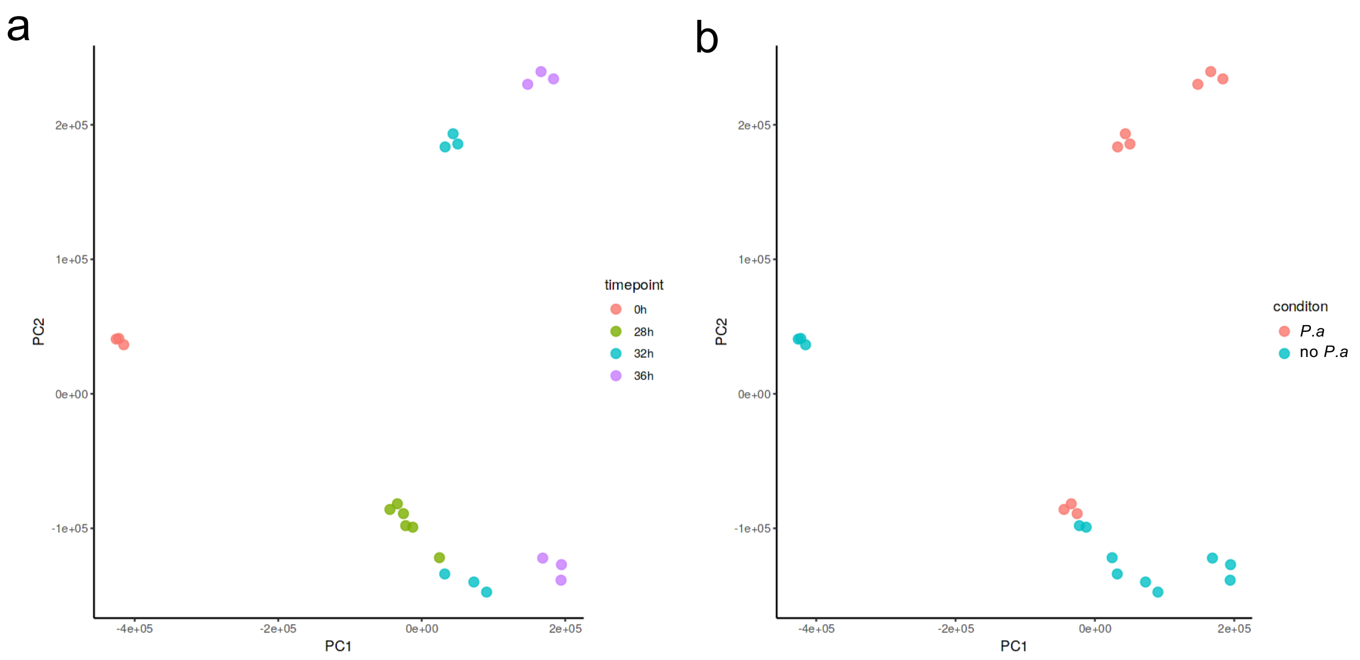


**Supplementary Fig. 3 |**

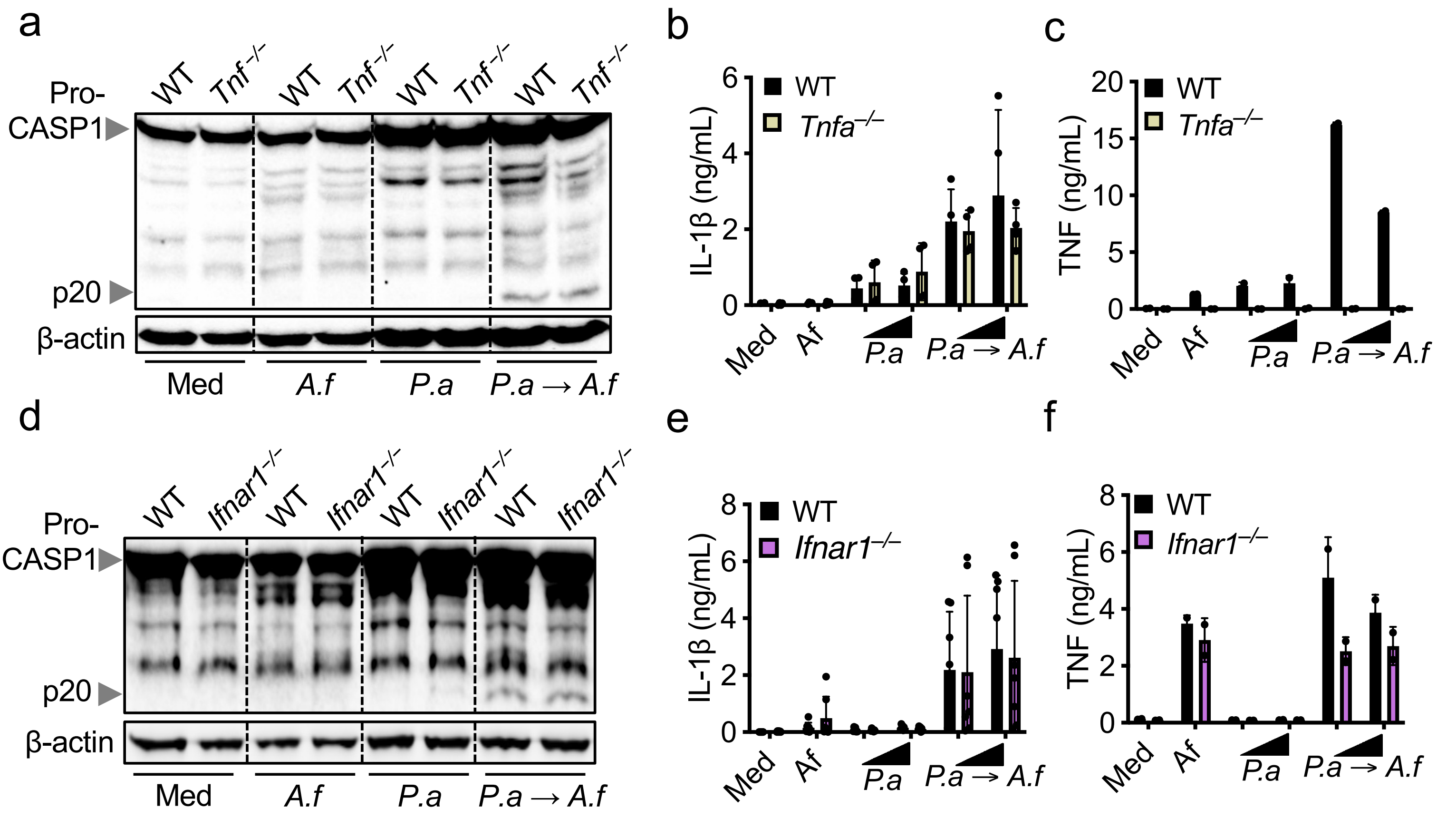


**Supplementary Fig. 4 |**

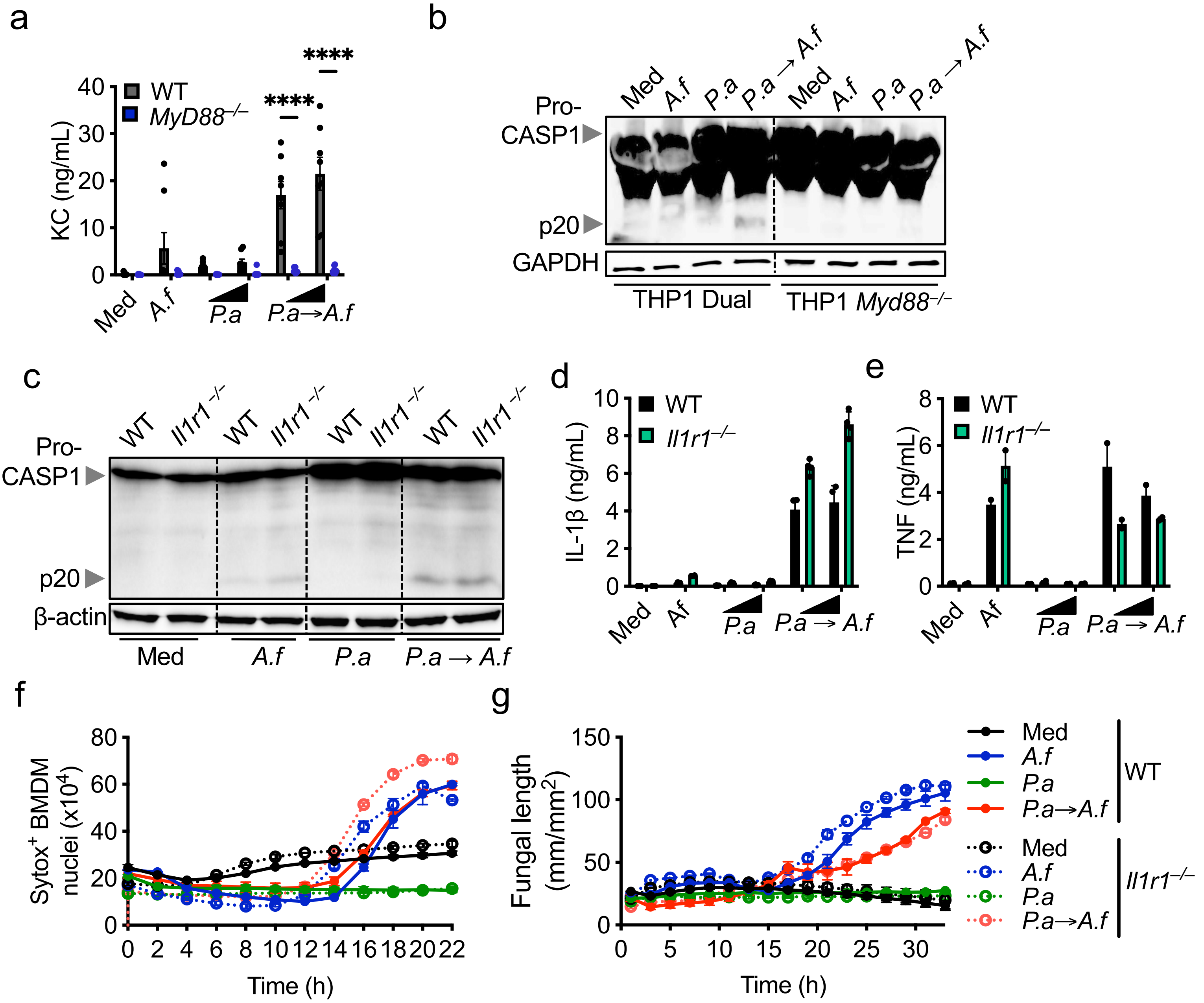


**Supplementary Fig. 5 |**

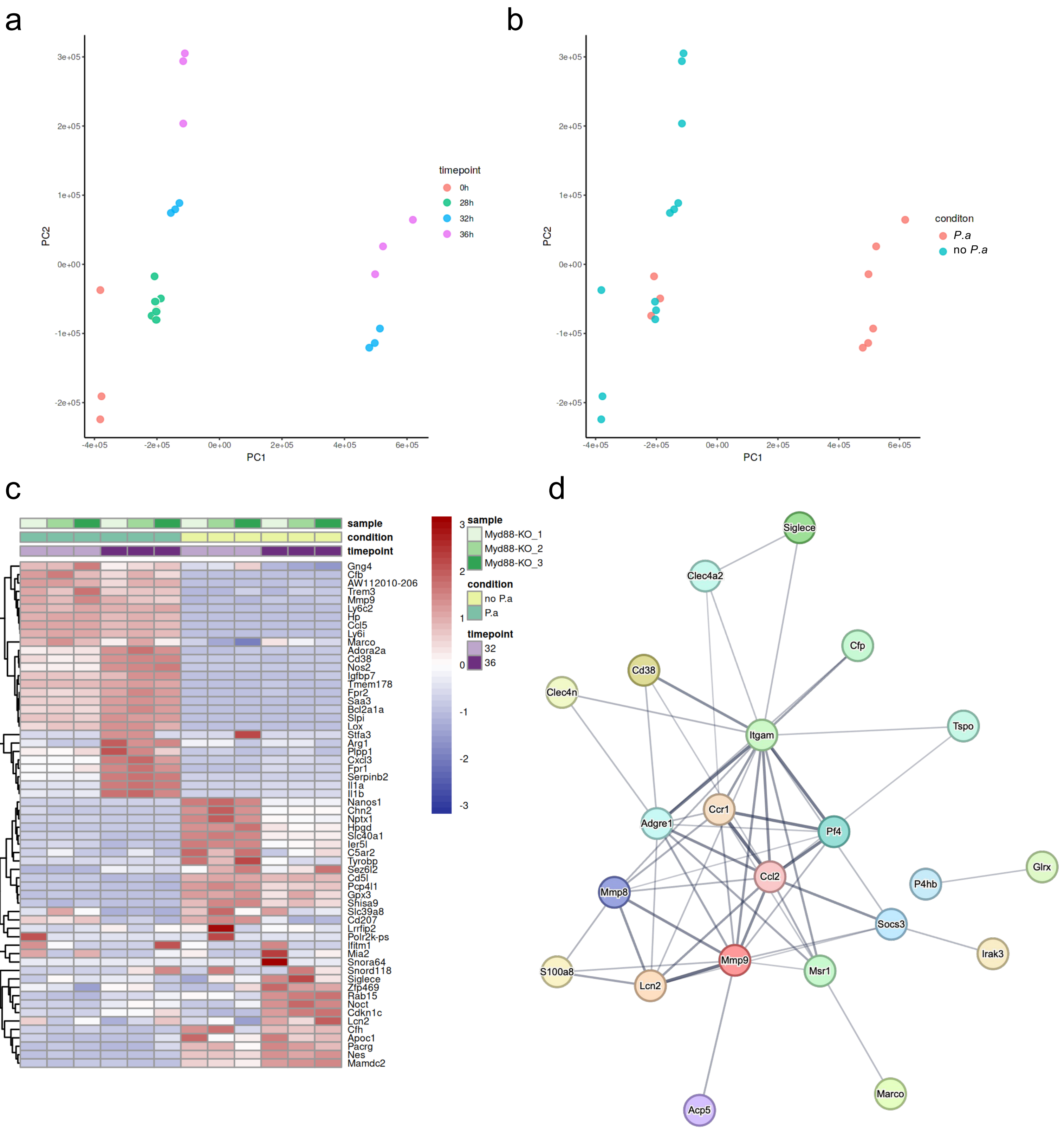


**Supplementary Fig. 6 |**

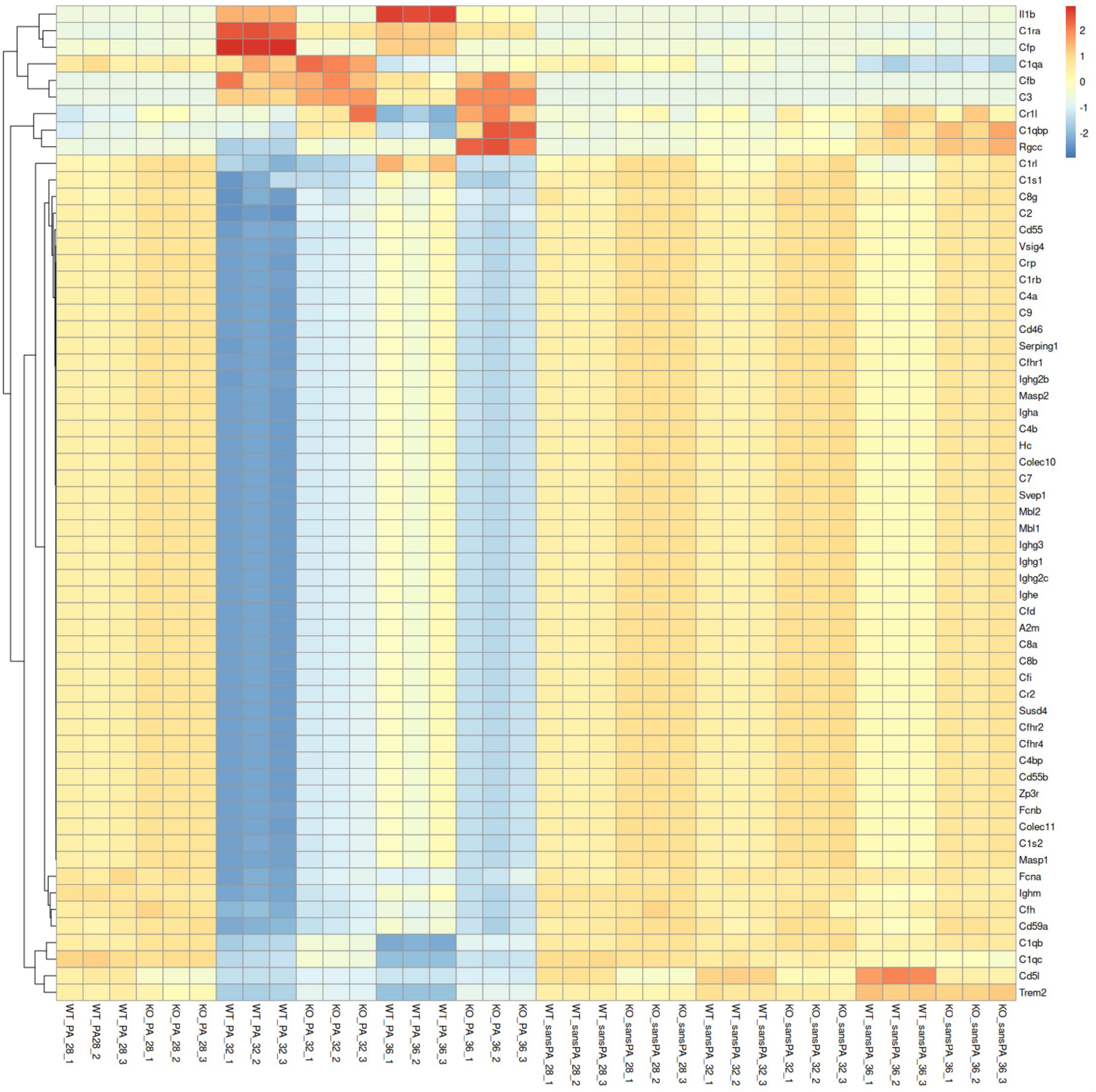


**Supplementary Fig. 7 |**

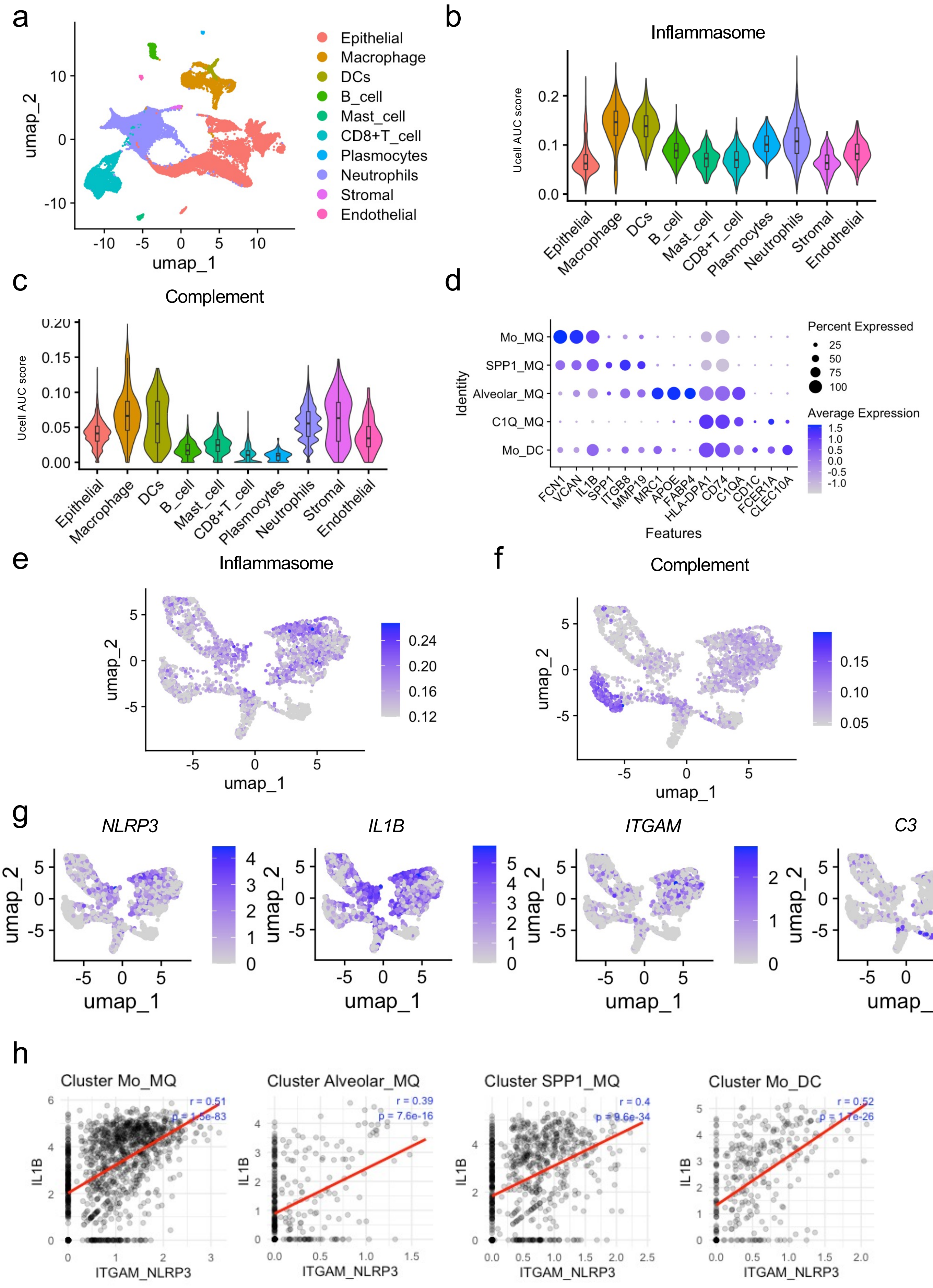


**Supplementary Fig. 8 |**

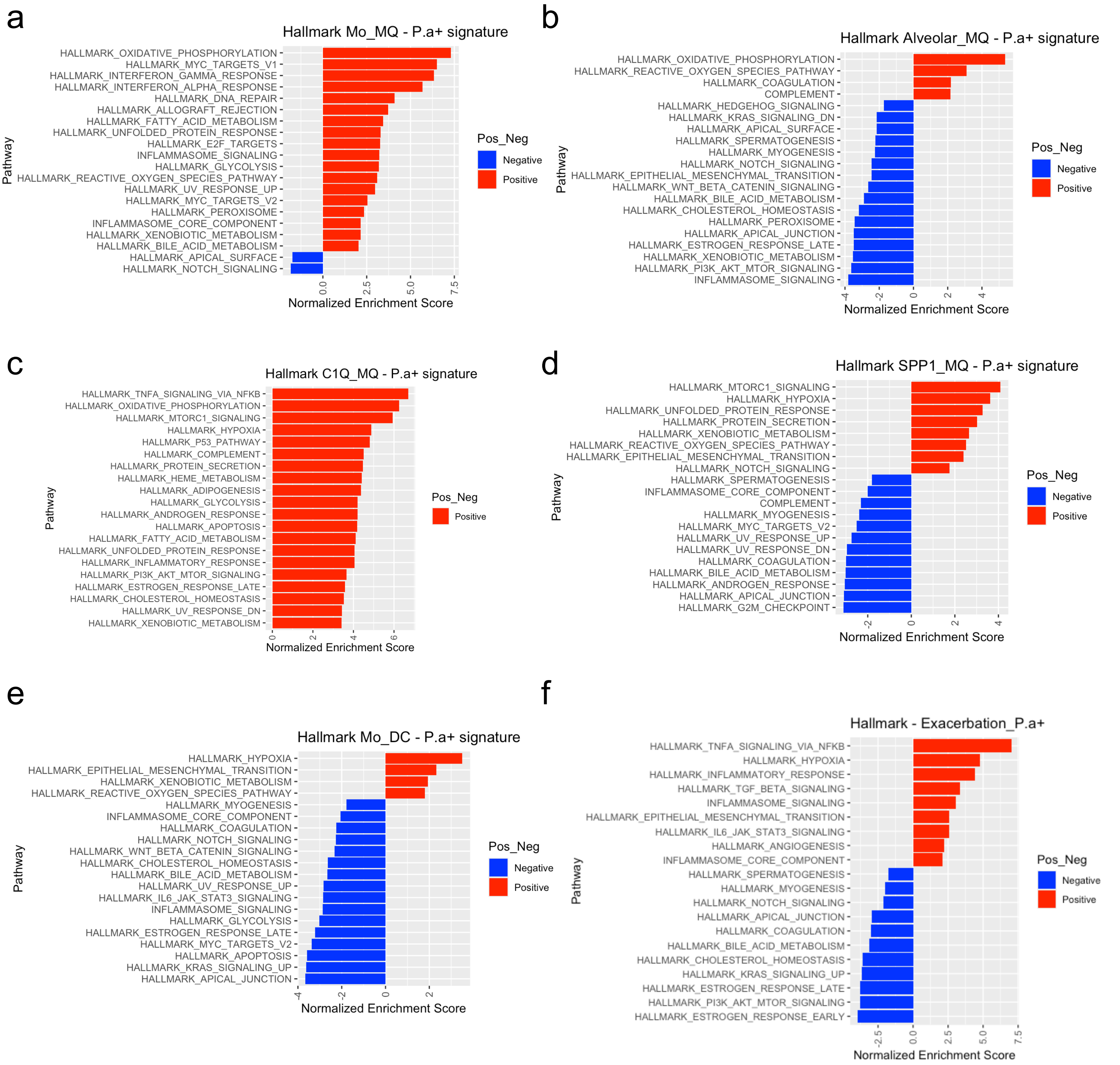


**Supplementary Fig. 9 |**

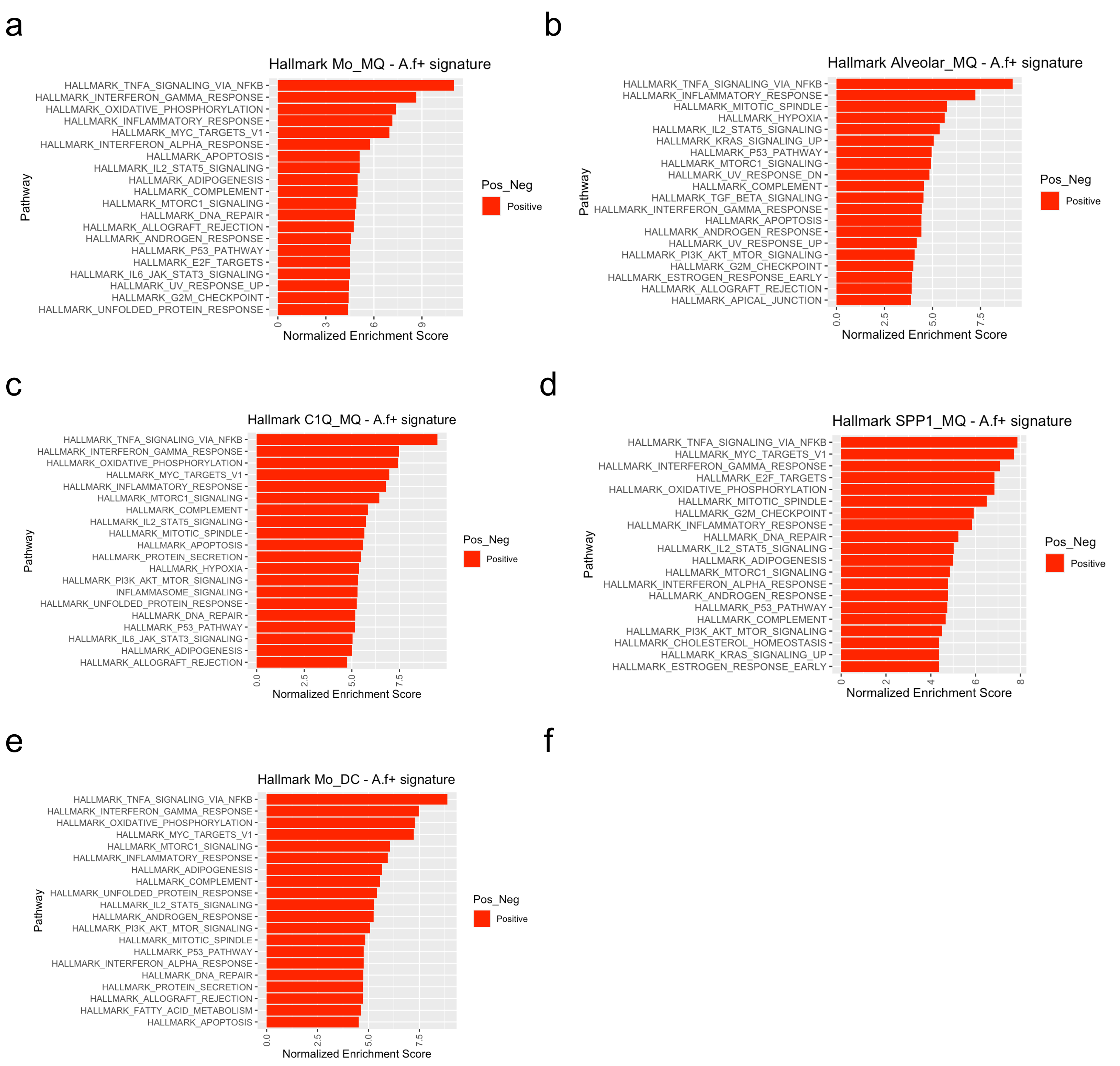


**Supplementary Fig. 10 |**

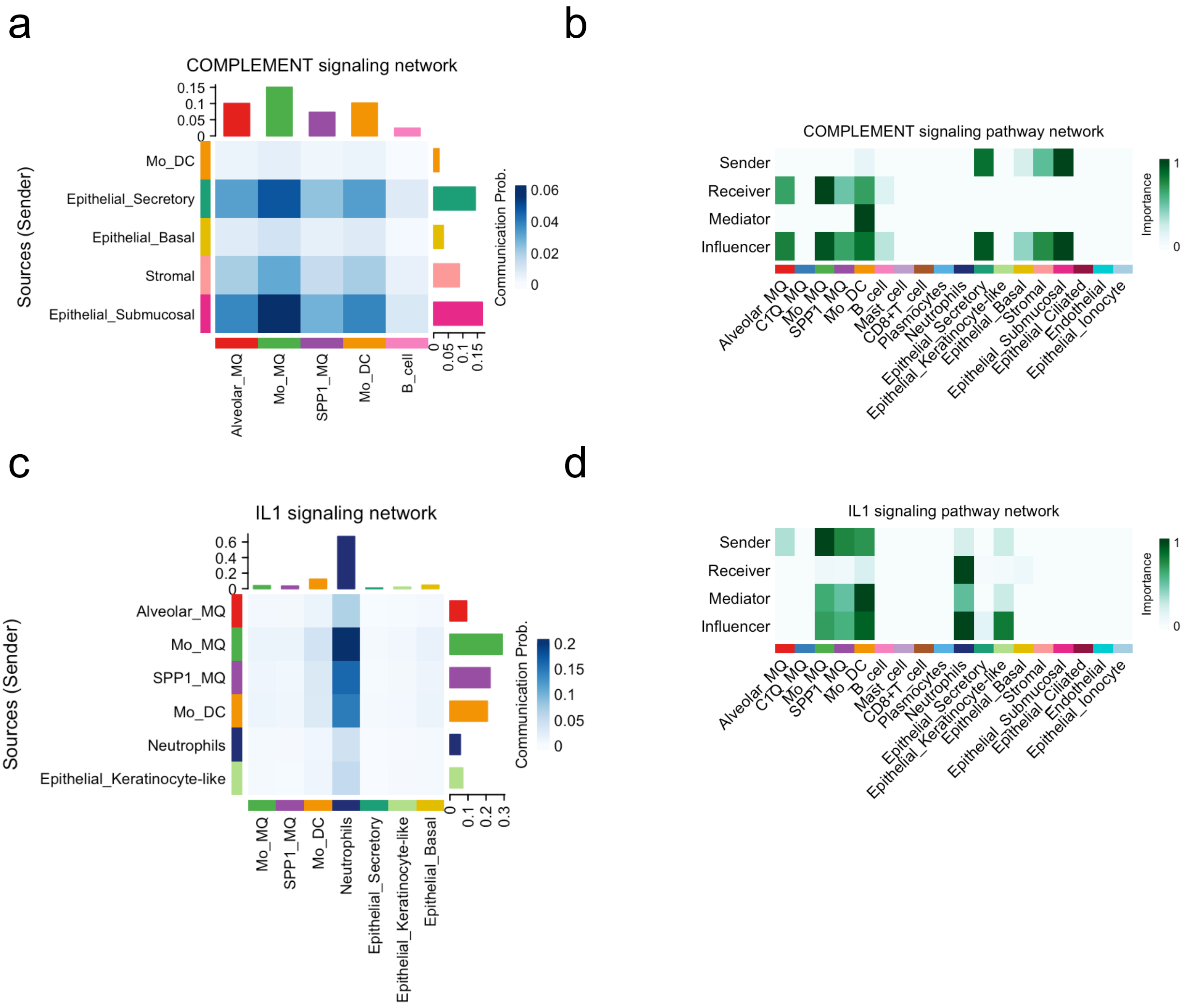


**Supplementary Fig. 11 |**
